# The temporal and genomic scale of selection following hybridization

**DOI:** 10.1101/2023.05.25.542345

**Authors:** Jeffrey Groh, Graham Coop

## Abstract

Genomic evidence supports an important role for selection in shaping patterns of introgression along the genome, but frameworks for understanding the dynamics underlying these patterns within hybrid populations have been lacking. Here, we develop methods based on the Wavelet Transform to understand the spatial genomic scale of local ancestry variation and its association with recombination rates. We present theory and use simulations to show how wavelet-based decompositions of ancestry variance along the genome and the correlation between ancestry and recombination reflect the joint effects of recombination, genetic drift, and genome-wide selection against introgressed alleles. Due to the clock-like effect of recombination in hybrids breaking up parental haplotypes, drift and selection produce predictable patterns of local ancestry variation at varying spatial genomic scales through time. Using wavelet approaches to identify the genomic scale of variance in ancestry and its correlates, we show that these methods can detect temporally localized effects of drift and selection. We apply these methods to previously published datasets from hybrid populations of swordtail fish (*Xiphophorus*) and baboons (*Papio*), and to inferred Neanderthal introgression in modern humans. Across systems, we find that upwards of 20% of the variation in local ancestry at the broadest genomic scales can be attributed to systematic selection against introgressed alleles, consistent with strong selection acting on early-generation hybrids. We also see signals of selection at fine genomic scales and much longer time scales. However, we show that our ability to confidently infer selection at fine scales is likely limited by inherent biases in current methods for estimating local ancestry from genomic similarity. Wavelet approaches will become widely applicable as genomic data from systems with introgression become increasingly available, and can help shed light on generalities of the genomic consequences of interspecific hybridization.

## 2 Introduction

The greater recognition in recent decades that introgression is a common feature of eukaryotic genomes has led to the view that species boundaries are semi-permeable (Harrison and Larson 2014). In this view, differential introgression along the genome is the result of selective filtering, with some neutral or widely favored alleles able to permeate into the genome of a hybridizing species, while others are restricted by genetic linkage to deleterious alleles in hybrids (Barton and Bengtsson 1986; Rieseberg *et al*. 1999; Wu 2001). While selection and assortative mating have long been thought to play an important role in maintaining species integrity in the face of gene flow, advancements in genomic sequencing and analysis have brought forth the possibility of reconstructing a more complete picture of genomic exchange between hybridizing species, forcing us to reckon with the vast complexity of how the genomic outcomes of hybridization are shaped by a dynamic interplay between recombination, selection, and genetic drift (Ravinet *et al*. 2017; Moran *et al*. 2021).

Following a hybridization event, recombination over multiple generations progressively breaks up contiguous segments of DNA inherited from the original source populations (ancestry tracts) into finer segments. Numerous genomic methods are now available to identify these tracts through genomic similarity to proxies for the source populations, and use the clock-like breakdown of tracts (or linkage disequilibrium between introgressed alleles) to make inferences about the timing of past mixture events (e.g. Sankararaman *et al*. 2008; Price *et al*. 2009; Maples *et al*. 2013; Loh *et al*. 2013). The changing spatial scale of co-inherited genetic material along the genome through time is simultaneously shaped by drift and selection acting at the population level, and will in turn influence how selection plays out in hybrid populations.

Genetic drift in a hybrid population shapes the ancestry proportion along the genome by increasing either ancestry state at random. Along the length of a chromosome, these deviations in the ancestry proportion will be autocorrelated due to the fact that an allele from one ancestry background that drifts to high frequency will tend to carry with it linked alleles from the same ancestry. For instance, if genetic drift is rapid during the early generations of hybridization, while ancestry tracts are long, broad contiguous portions of the genome might randomly fix for either ancestry. Conversely, in a large population where genetic drift is slow, by the time an allele of one ancestry reaches fixation, it will have been unlinked from all but the closest neighboring alleles from the same source population. The progressive shortening of ancestry tracts is slowed and ultimately stopped by genetic drift; once a genomic segment fixes for either ancestry state, recombination events within the segment will cease to create ancestry breakpoints in the descendant chromosomes (Gravel 2012; Liang and Nielsen 2014).

Selection acting on hybrids will further shape variation in levels of introgressed ancestry along the genome. Hybrids often experience strong selection, as their genomes can contain genetic incompatibilities or encode maladaptive phenotypes (Coyne and Orr 2004). Selection for or against an introgressed allele in an admixed population will lead to an excess or depletion of the corresponding ancestry in the surrounding region, the length of which depends on the strength and timing of selection relative to the timing of admixture (Maynard Smith and Haigh 1974). This concept has been leveraged to identify selected loci in recently admixed populations (Tang *et al*. 2007; Hamid *et al*. 2021) and date the onset of selection on introgressed alleles (Yair *et al*. 2021). Similarly, strong selection against introgressed alleles in specific genomic regions is thought to have contributed to the formation of so-called ‘introgression deserts’ (Sankararaman *et al*. 2014; Hajdinjak *et al*. 2021). Analogously, variable patterns of divergence between species along the genome are thought to form at least in part through barriers to gene flow, i.e. selection preferentially removing introgressed haplotypes in regions harboring incompatibilities or loci contributing to divergent adaptation (Nosil *et al*. 2009; Berner and Roesti 2017; Samuk *et al*. 2017; Aeschbacher *et al*. 2017).

Increasing attention has focused on forming a more general understanding of how the distribution and frequency of introgressed ancestry along the genome has been shaped by natural selection (Sankararaman *et al*. 2014; Juric *et al*. 2016). Various studies have identified genome-wide correlations between minor parent ancestry proportion and recombination rate, indicating that selection has acted at many loci throughout the genome to remove alleles from the minor parent ancestry (Schumer *et al*. 2018; Martin *et al*. 2019; Edelman *et al*. 2019; Calfee *et al*. 2021; Vilgalys *et al*. 2022). Such correlations emerge due to the slower decay of linkage disequilibrium (LD) between deleterious alleles carried on introgressed segments in low recombination regions, allowing for selection to more efficiently remove linked introgressed alleles in these regions (Nachman and Payseur 2012; Veller *et al*. 2023). Whereas these correlations can represent a snapshot of the cumulative effects of selection over tens to thousands of generations, theoretical work has shown that the strength of selection acting on hybrids likely varies dramatically through time. Due to extensive admixture LD in early-generation hybrids, the selective effects of many introgressed alleles combine, creating very strong selection on individuals carrying introgressed haplotypes (Barton 1983). Thus, under a model of selection against introgressed alleles at many loci throughout the genome, the rate of removal of introgressed ancestry is greatest in the first several generations following hybridization (Harris and Nielsen 2016; Veller *et al*. 2023).

We now have clear genomic evidence that selection plays a role in maintaining species in the face of hybridization, but thus far have lacked a methodology to disentangle the temporal effects of selection, and to understand how selection shapes spatial ancestry patterns in the genome. Here, we develop genomic methods for analyzing temporal dynamics of drift and selection in hybrid populations based on the Discrete Wavelet Transform (DWT), a tool commonly used in time series analysis. After introducing important features of the DWT, we show how the ancestry variance present at different genomic scales can be captured by the wavelet variance decomposition, and how this captures the time scale of evolutionary processes. We further show that a wavelet decomposition of the correlation between introgressed ancestry proportion and recombination rate, which decomposes the correlation into contributions of different scales, tracks temporal dynamics of genome-wide selection acting against one ancestry. Finally, we apply these methods to three empirical data sets: time-series data from a hybrid population of swordtail fish (*Xiphophorus*), a hybrid swarm between yellow and anubis baboons (*Papio*), and Neanderthal ancestry in modern humans. Across all data sets, we find patterns consistent with selection beginning early after hybridization and continuing throughout multiple generations.

### 2.1 Spatial genomic decomposition of ancestry variance using the Discrete Wavelet Transform

A common goal in the analysis of temporal or spatial signals is to understand the scale of variation present in the signal (e.g. using methods such as the Fourier decomposition). One approach is the wavelet transform, which can be used to understand the scale of variation in a signal as well as capture information about local signal features. Although widely used in physical sciences, the wavelet decomposition has seen more limited application in population genomics (but see Spencer *et al*. 2006; Pugach *et al*. 2011; Chan *et al*. 2012; Sanderson *et al*. 2015, for examples).

We start with a signal of interest, *x*(𝓁), such as ancestry state *x* measured at a set of evenly-spaced locations 𝓁 = 1, …, *L* along a contiguous chromosome of length *L*. If the data are not evenly spaced, we first interpolate to obtain evenly-spaced measurements, along either a genetic or physical map of a chromosome. We will use the Discrete Wavelet Transform to decompose the variation in this signal - that is, the variation in the values of the signal measured along the chromosome - into components associated with a discrete set of spatial genomic scales. This is accomplished through multiplying our signal with a set of wavelets, functions written as *ψ*_*λ,i*_(𝓁) that capture changes in the signal over varying spatial scales and locations. Informally, each wavelet resembles a finite wave, oscillating equally between positive and negative values over some characteristic scale *λ* centered on some location *i*. While many such functions exist, we use Haar wavelets (examples shown as black lines in Fig. 1A), which take a negative constant value for a stretch of sequence of length *λ*, switching to a positive constant value at location *i*, and are zero everywhere else.

**Figure 1:**
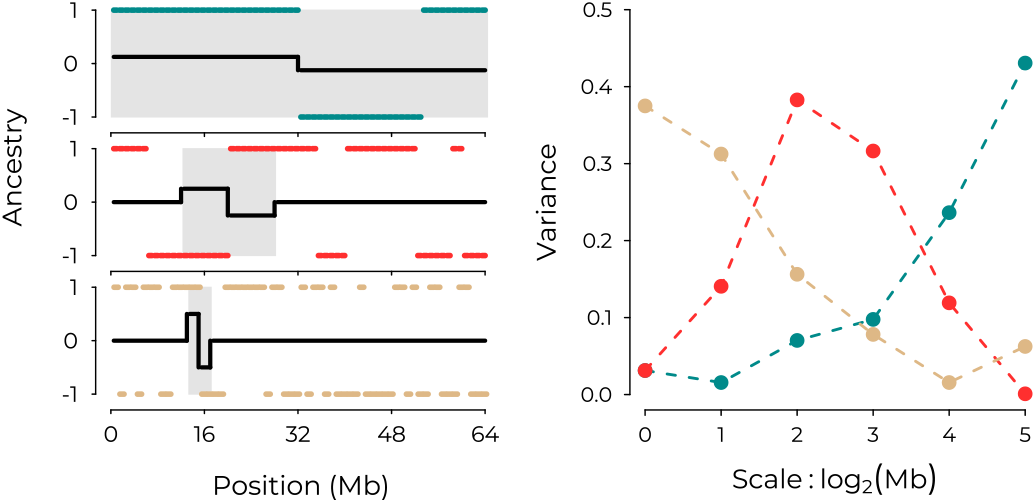
**(Left)** Ancestry states *x*, coded arbitrarily as *x*(𝓁) = {1, 1} along three separate chromosomes, with examples of Haar wavelets with different scales and positions overlaid in black. From top to bottom, ancestry tracts are shorter, representing different histories of recombination. Shaded intervals highlight the portion of the ancestry signal contributing to the resulting wavelet coefficient. (Top left) Positive covariance between a *ψ*_*λ*=6_ wavelet and *x* within the shaded interval yields a positive wavelet coefficient corresponding to a change in average ancestry state over the two halves of the chromosome. (Middle left) Negative covariance between a *ψ*_*λ*=4_ wavelet and *x* within the shaded interval yields a negative wavelet coefficient. (Bottom left) Positive covariance between a *ψ*_*λ*=2_ wavelet and *x* within the shaded interval gives a positive wavelet coefficient. **(Right)** The complete set of squared wavelet coefficients determines the power spectrum for the three ancestry signals, correspondence indicated in color.

The DWT transforms our signal into a set of wavelet coefficients, *w*_*λ,i*_, each of which measures the strength of association between the signal and a corresponding wavelet:

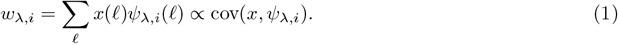

Because the mean of a wavelet is zero, a wavelet coefficient is proportional to the covariance between the signal and the corresponding wavelet, and thus measures the extent to which the wavelet captures variation in the signal over scale *λ* and location *i*. In the case of Haar wavelets, each wavelet coefficient measures a deviation between two adjacent windowed averages of the signal for a specific window size and location.

The set of wavelet coefficients produced by the DWT retains all of the information in the original signal while avoiding redundancy. To achieve this, the wavelet scales *λ* are chosen to form a doubling series (e.g. 1kb, 2kb, 4kb, 8kb, …), and any two wavelets of different scale have zero covariance. Thus, variation in the signal at each scale is measured independently from variation measured at any other scale. This property distinguishes our approach from window-based approaches commonly used in genomics research, where statistics calculated from genomic windows of varying sizes are confounded, due to the fact that smaller windows are nested within larger windows.

The wavelets at a given scale are simply shifted versions of each other, with their non-zero portions covering different portions of the sequence. In the traditional DWT, these are placed such their non-zero portions cover the entire sequence without any overlap. Thus, any two wavelets of the same scale also have zero covariance, avoiding redundancy between neighboring wavelet coefficients. We instead use a modified and more flexible version of the traditional DWT known as the MODWT, which, for a given scale, uses wavelets placed at all positions in the sequence (Percival and Walden 2000). This yields less noisy estimates of the wavelet variance and covariance (described below) and retains the property of measuring variation independently across scales. Further details on the wavelet transform are given in Appendix A.

#### Wavelet variance decomposition

The average of the squared wavelet coefficients at a given scale, called a wavelet variance, has the interpretation of the variance in our signal associated with *changes* in the value of the signal occurring at that scale. The total variance of the original signal along the sequence, 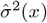, can be decomposed as the sum of wavelet variances across scales, also known as the power spectrum:

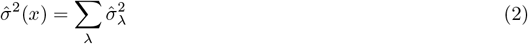

Since the wavelet scales *λ* are powers of two, if our sequence length is not itself a power of two, then there will be an additional component of leftover variance - referred to in the wavelet literature as scaling variance - due to the largest-scale wavelet not covering the entire sequence. As we average across chromosomes of different lengths, we fix the resolution of measurement (e.g. 50kb or 2^−10^ Morgans) such that L is not a power of two, and so we are left with scaling variance. To simplify interpretation, we omit this scaling variance from the results shown in the main text as it represents only a minor component of the total variance in all our analyses.

When applied to ancestry state of a haploid copy of a chromosome, the power spectrum provides a summary of the length distribution of ancestry tracts (Fig. 1), with long ancestry tracts generating variance at broad scales (blue lines) and shorter ancestry tracts generating variance at finer scales (tan lines). This property is leveraged in the wavelet-based admixture dating methods of Pugach *et al*. (2011) and Sanderson *et al*. (2015). While recombination is the primary force determining the lengths of admixture tracts for single chromosomes, genetic drift and natural selection acting in a hybrid population will cause homologous pairs of the same chromosome within a population to co-vary in their ancestry state. The spatial extent of this covariance is captured by applying the power spectrum to mean ancestry across multiple segregating copies of the same chromosome, i.e. the ancestry proportion (Eqn. S15).

To obtain a complete spatial decomposition of the variance in ancestry proportion along the whole genome, we apply the wavelet transform separately to each chromosome (e.g. the ancestry proportion along chromosomes 1, 2, …, N) and take a chromosome length-weighted average of wavelet variances across chromosomes at each scale. (Due to heterogeneity in chromosome sizes, variation at the largest scales is present only on some chromosomes, and so we estimate wavelet variances using only the chromosomes for which a given scale is present.) Finally, since wavelet variances give only the within-chromosome portion of ancestry variance representing fluctuations around the mean for each chromosome, we account for the among-chromosome variance contribution by calculating a weighted variance of chromosome means of ancestry. This among-chromosome variance will be labelled *chrom* in the results. We can separately calculate the proportion of total genomic variance explained by each component; if all chromosomes have the same length, these quantities will be the same up to a constant. These components combined (wavelet variance, scaling variance, and among-chromosome variance) form a complete variance decomposition of our measured ancestry signal across the genome.

#### Wavelet correlation decomposition

Wavelet methods can also be applied to examine the scale of covariation between two signals (e.g. ancestry state and recombination rate), which we will use to examine temporal dynamics of selection. The overall correlation between two signals *x* and *y* (measured at the finest resolution available) can be decomposed into the sum of contributions from each scale:

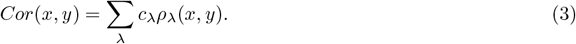

where the correlation *ρ*_*λ*_(*x, y*) between our two signals at scale *λ* is weighted by an average proportion of variance explained by scale *λ* in the two signals, *c*_*λ*_ (see e.g. Spencer *et al*. 2006, Text S2). The correlation at scale *λ* is computed from the wavelet coefficients of *x* and *y* at scale *λ* and measures the strength of association between localized directional changes in *x* and *y* around their mean value at that scale. As with the variance decomposition, a complete decomposition of the overall genome-wide correlation will include additional terms due to leftover portions of chromosome not covered by the largest wavelets, as well as an among-chromosome component.

## 3 Results

### 3.1 The wavelet variance captures temporal effects of genetic drift in hybrid populations

We first illustrate the effects of genetic drift on the wavelet variance decomposition of ancestry state in a hybrid population in the absence of selection. Following a pulse of hybridization as recombination shortens ancestry tracts, genetic drift meanwhile generates deviations in the ancestry proportion along the genome away from the initial mixture proportion. The timescale of drift relative to recombination in hybrids determines the spatial scale of auto-correlation in these deviations, that is, the spatial scale of variance in mean ancestry along the genome. If a genetic bottleneck occurs shortly after the mixture event, the large deviations in ancestry proportion it causes will happen while ancestry tracts are long, so these deviations will be auto-correlated over broad scales (maroon, Fig. 2, top panel). In contrast, drift in a large constant-sized population generates comparable ancestry deviations over much longer timescales (blue, Fig. 2, top panel), by which point recombination has had more time to whittle down ancestry tracts, such that the variance in mean ancestry along the genome due to drift is on finer spatial scales.

**Figure 2:**
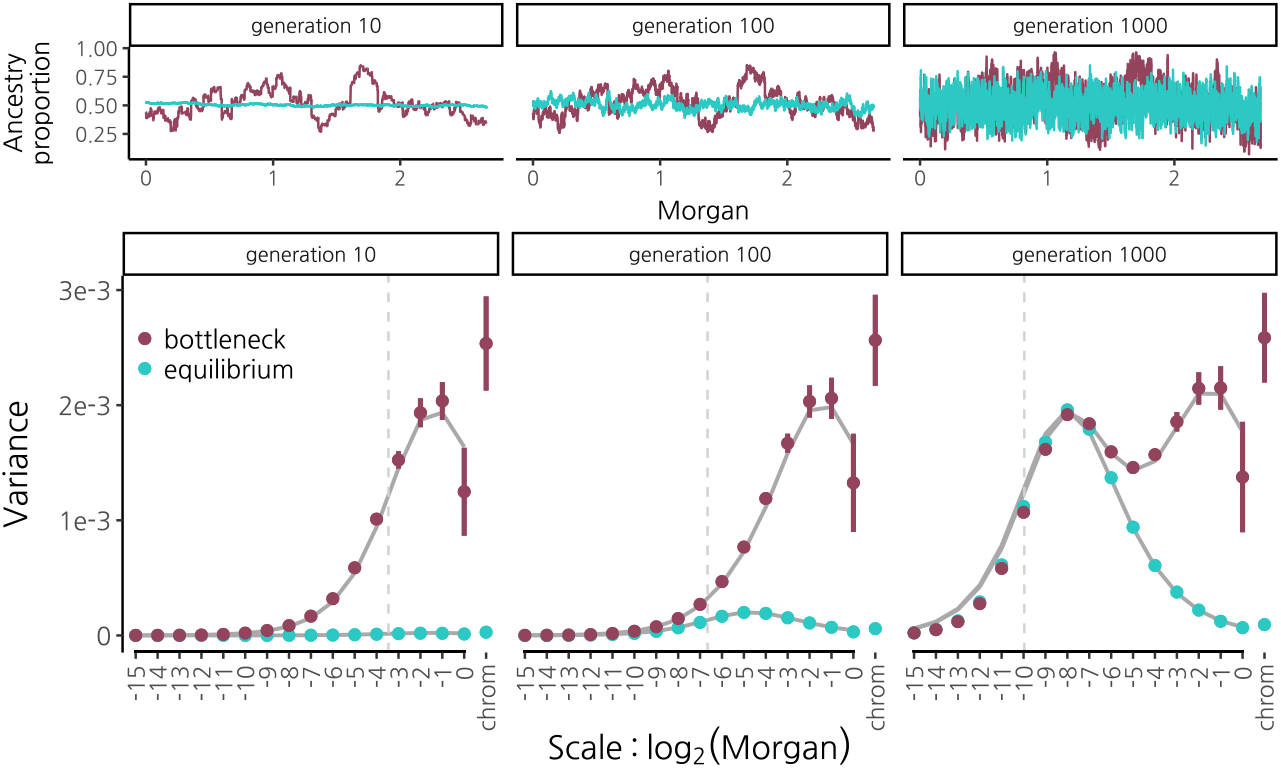
Wavelet variance decomposition through time of ancestry proportion in a 50/50 population mixture undergoing genetic drift with recombination and no selection. We simulated (using SLiM, Haller and Messer, 2019) a population of constant size 2N=20000 (blue) and a population that undergoes a bottleneck to 2N=200 for just the first 10 generations of recombination in hybrids, then expands to 2N=20000 (maroon). **(Top)** Ancestry proportion along human chromosome 1 from a single simulation run. From left to right, shown after 10, 100, and 1000 generations of recombination in hybrids. **(Bottom)** Wavelet variance decomposition showing the spatial scale of variance in ancestry proportion. Points and error bars show means and 95% confidence intervals across 20 replicate simulations. Solid grey lines show theoretical expectations. Vertical dotted grey lines indicate the expected distance between recombination breakpoints that have accrued along a single chromosome since the hybridization pulse. Note that since results are shown on the genetic map, recombination rate variation does not influence these patterns, other than through interpolation error.

The wavelet power spectrum of mean ancestry provides an elegant summary that captures these effects. A strong bottleneck concurrent with a hybridization pulse generates large wavelet variances at broad scales (maroon, Fig. 2 bottom panel). Importantly, this broad-scale variance is maintained through time; even after 1000 generations, these large wavelet variances at broad scales retain the signature of the early bottleneck. So long as ancestry remains polymorphic at many loci, genetic drift continues to generate variance in mean ancestry at progressively finer scales through time. Thus, following the bottleneck and population expansion, wavelet variances build at finer scales according to the rate of drift in the larger population. Contrast this to the case of a large, constant-sized population where variance along the genome only starts to become apparent at fine scales many generations after mixture (blue, Fig. 2 bottom panel).

We derived the expectation for the wavelet variance using coalescent theory (Appendix B), and find that it shows good agreement with simulation results (Fig. 2, solid grey lines in bottom panel). We note however that subtle biases in our simulated wavelet variances (e.g. downward bias at fine scales in generation 1000) result from the interpolation of ancestry state between simulated loci that are evenly spaced on a physical map to locations that are evenly spaced on a genetic map (Fig. S2).

### 3.2 Measuring the timescale of selection on introgressed ancestry

As with drift, selection generates deviations in mean ancestry along the genome away from the initial mixture proportion, with the extent of auto-correlation in these deviations determined by the timing and strength of selection relative to the timing of mixture, as well as variation in recombination rate along the sequence. To explore the role of selection in shaping ancestry variation, we performed forward simulations of a hybridization pulse followed by selection acting additively against alleles fixed in one ancestry background at many loci (10,000) genome-wide on a genetic map modelled on the human autosomes (for example representing selection due to polygenic adaptation of one source population to the local environment). We chose this highly polygenic model in part to provide fine-scale variation for selection to act upon, but also discuss models with selection on fewer loci below.

We find that selection acting to remove introgressed alleles at multiple loci distorts the power spectrum towards proportionally greater ancestry variance at broad scales relative to the neutral expectation (Fig. 3A). This effect is seen across a range of numbers of loci (10 - 10,000) under selection in hybrids, with greater broad-scale variance generated when the same total additive selection strength is distributed across fewer loci (Fig. S4). Separately, selection can decrease levels of ancestry variance across some or all scales (depending on the recombination map) relative to the neutral expectation by virtue of moving the introgressed ancestry proportion towards to zero (Fig. S5). Note that widespread weak selection may lead to departures from the neutral power spectrum on the same order as those caused by interpolation. Thus, in comparing empirical results to neutral expectations it may in some cases be preferable to use simulations that incorporate the interpolation noise.

**Figure 3:**
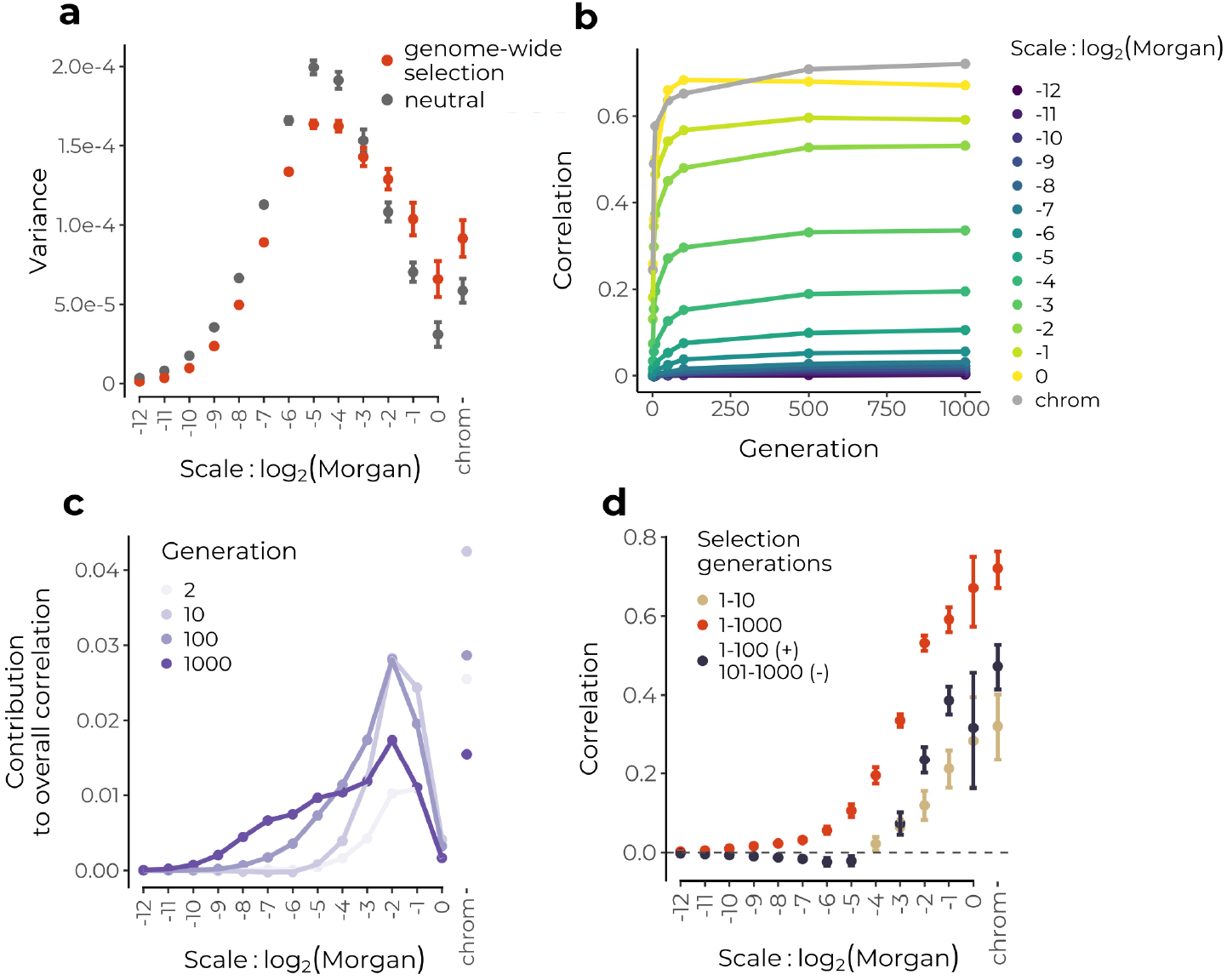
Simulations of selection following a pulse of hybridization starting from a 50/50 population mixture.**(a)** Genome-wide selection in conjunction with broad-scale variation in recombination rates leads to differential removal of introgressed ancestry at broad scales, thereby biasing the power spectrum towards greater variance at these scales compared to the neutral expectation (viewed after 100 generations). **(b)** Selection against many alleles on one ancestry background rapidly establishes broad-scale correlation between recombination and minor parent ancestry in early generation hybrids. **(c)** The overall correlation is dominated by broad-scales, but finer scales contribute increasingly more through time. **(d)** Selection acting only on the first 10 generations of recombinant hybrids generates significant positive wavelet correlations only at broad scales (brown) (viewed in generation 1000), whereas continuous selection over 1000 generations continues to generate correlations on finer scales (red). When selection acts continuously but reverses direction after 100 generations to favor the alternate ancestry, positive broad-scale correlations persist as negative correlations establish at finer scales. Only significant correlations are shown, error bars represent 95% confidence intervals across 20 replicate simulations.

Critically, variance in mean ancestry along the genome produced by systematic selection against one ancestry can be distinguished from that produced by genetic drift; we expect introgressed ancestry depletions against the genome-wide background to be spatially associated with features that govern the strength of selection, namely the recombination rate and the density of selected sites. Indeed, positive correlations between minor parent ancestry and recombination rate or negative correlations with coding density are often the basis for inferring genome-wide selection against alleles from one ancestry (e.g. Schumer *et al*. 2018).

As selection over multiple generations establishes these correlations against a shifting backdrop of the spatial scale of ancestry variance due to recombination, the wavelet spatial decomposition of the correlation between ancestry state and recombination (Eqn. 3) can be used to track the effects of selection through time. In the early generations after hybridization, when selection sees numerous selected alleles linked in long ancestry tracts, strong correlations rapidly establish at the broadest genomic scales (Fig. 3B). Correspondingly, as the bulk of variance in mean ancestry at this stage is also at broad scales, these scales contribute most to the total correlation (Fig. 3C). Wavelet correlations at finer scales increase more gradually through time as selection continues to operate on an increasingly fine-scale mosaic of ancestry tracts. As drift increases variance at finer scales and selection generates correlations on these scales (selection may be increasing or decreasing these variances, Fig. S5), fine scale correlations contribute more the overall correlation through time (Fig. 3C). In simulations where the same strength of selection is spread over fewer loci under selection in hybrids, we find that correlations are weaker and do not continue to establish at finer scales through time (Fig. S6). This makes intuitive sense, as we only expect selection to creates fine-scale correlations with recombination if selected loci are distributed over fine scales. In each cases, broad scales continue to constitute a significant portion of the overall correlation through time, indicating that early selection has an outsized influence on overall patterns of ancestry in hybrids.

As we observed that the power spectrum of mean ancestry could preserve a memory of an early bottleneck after hybridization, we next asked whether the wavelet correlation decomposition could similarly be used to detect temporally-localized effects of selection. We thus simulated a scenario where selection acted only on the first 10 generations of recombinant hybrids after admixture. In that case, correlations between recombination and ancestry proportion remained restricted to only the largest scales and remained present after 1000 generations of recombination in hybrids (Fig. 3D, brown). This suggests that observing significant fine-scale wavelet correlations between recombination and ancestry proportion on the genetic map indicates that selection continues to have wide-spread effects on ancestry in later generations of mixture. We next simulated two additional scenarios, where (1) selection begins only after 500 generations of neutral mixture, and (2) where selection acts in every generation following the hybridization pulse but switches directions after 100 generations to favor the alternate ancestry allele at each locus (these model were chosen for illustrative purposes and not to necessarily reflect biologically realistic scenarios). We find that while selection acting only in later generations also readily generates significant broad-scale correlations with recombination (scenario 1), broad-scale correlations that are established by early selection are not reversed even by subsequent generations of selection acting in the opposing direction (scenario 2) (Figs. 3D, S7). Thus, the wavelet decomposition of the correlation between mean ancestry and recombination is capable of revealing disparate effects of selection in different time periods.

In the analyses above, we modelled a single pulse of admixture to provide intuition for how the wavelet decompositions capture temporal dynamics of drift and selection on introgressed ancestry. In reality, hybrid populations may receive multiple influxes of new parental individuals, or exist as a hybrid zone between larger populations. Under such models, the lengths of introgressed segments will have a mixture distribution reflecting the cumulative effects of multiple hybridization events, and the resulting wavelet decompositions will capture the effects of selection and drift on this combined distribution of segments. While we demonstrated the wavelet decompositions using a specific model of demography and selection, the methods themselves are agnostic to any assumptions about the underlying model of hybridization.

We next apply the wavelet methods illustrated above to previously published ancestry calls from several empirical datasets. In our theoretical and simulation work, ‘ancestry’ can be defined precisely, as we directly track the descent of haplotypes from either of two well-defined populations that form a mixture at a specified time in the past. In reality, this information is not known and may be poorly defined, and ancestry for hybrid (or admixed) populations is defined with respect to genetic similarity to sets of reference samples (A and B) that are thought to be representative of the variation present in the original mixing groups. Thus, in describing specific analyses we use terminology such as A-like haplotypes to refer to regions of the genome that have been computationally identified as more similar to reference sample A than sample B (Coop 2022; National Academies of Sciences, Engineering, and Medicine 2023). We use the term ancestry when we discuss the inferences we draw from these analyses, e.g. that selection acts against alleles from the species A ancestry, reflecting the fact that our inferences are placed in a conceptual model of two divergent populations mixing upon secondary contact.

### 3.3 Application to hybrid swordtail fishes

To examine the roles of recombination, drift and selection in shaping ancestry patterns in a recently-formed hybrid population, we analyzed time-series whole genome data from a hybrid population of swordtail fish from Acuapa river in Hidalgo, Mexico. This population is thought to have formed from hybridization between *Xiphophorus birchmanni* and *X. malinche*, within the last 100 years (Powell *et al*. 2021). In several independently-formed hybrid zones between the same species, Schumer *et al*. (2018) observed positive correlations between recombination rate and minor parent ancestry proportion, implicating selection in shaping ancestry patterns genome-wide. Furthermore, numerous incompatibilities are known to segregate in these hybrid populations (Powell *et al*. 2020; Moran *et al*. 2022).

We made use of previously-inferred local ancestry patterns in a set of temporally staggered samples ranging from 2006 to 2018 (Powell *et al*. 2021). Genotype posterior probabilities of matching allopatric *X. birchmanni* / *X. malinche* alleles were called using a Hidden Markov Model (HMM) at a set of loci along the genome that are highly differentiated between the species, and we interpolated the average marginal posterior probability of matching the *X. malinche* allele to yield the sample proportion of malinche-like haplotypes at evenly-spaced intervals along the genome.

To illustrate the effects of recombination across the sampling interval, we visualized the power spectrum of the proportion of malinche-like haplotypes on the genetic map for each collection year. As expected by the shortening of ancestry tracts through time, the power spectrum shifts towards greater variance at finer scales across the temporal sampling interval (Fig. 4A). Increasing levels of overall variance in the signal through time are not due to differences in sample size and are consistent with expected effects of genetic drift following hybridization (Fig. S8, and may also reflect selection).

**Figure 4:**
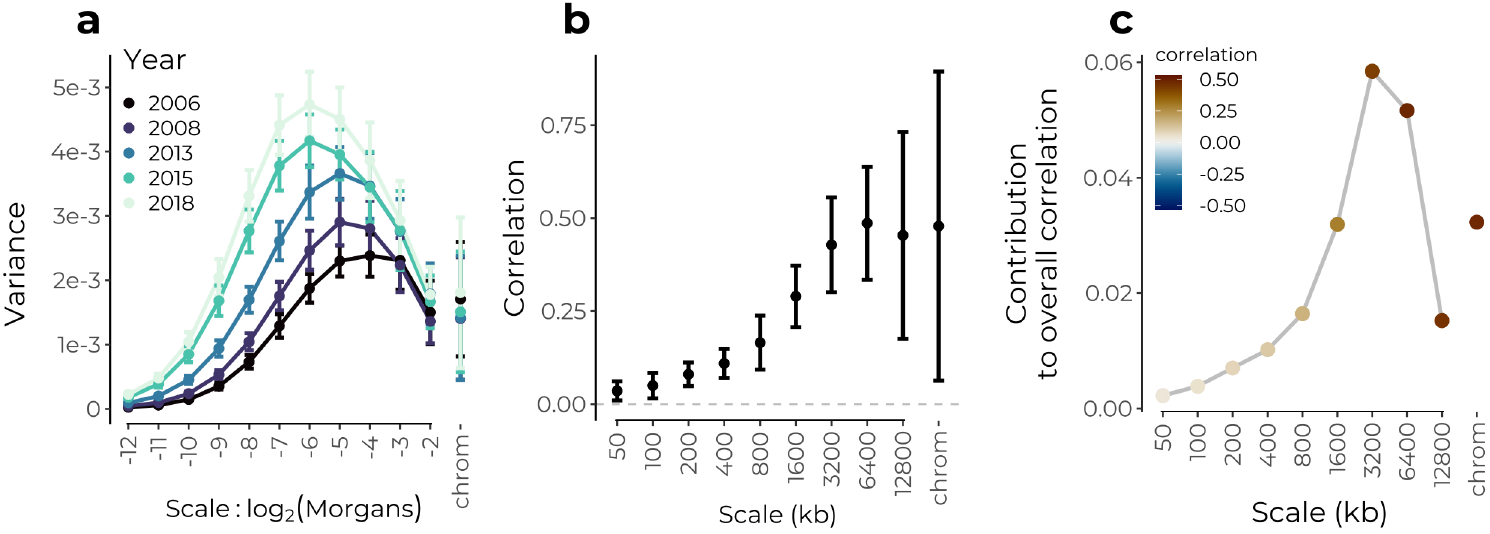
Wavelet analysis of hybrid genomes in a population of swordtail fish from Acuapa River in Hidalgo, Mexico. **(a)** Power spectrum of the proportion of malinche-like haplotypes on the genetic map for five time points points between 2006 and 2018. Wavelet variances represent a weighted averaged across chromosomes, and error bars are 95% confidence intervals from a weighted jackknife of chromosomes. **(b)** Correlations at each spatial genomic scale (on the physical map) between wavelet coefficients for the proportion of malinche-like haplotypes and wavelet coefficients for recombination rate. Squared values give an estimate of the proportion of variance in ancestry state explained by systematic selection against malinche-like alleles. Data shown only for 2006 sample, patterns similar across years. **(c)** Contribution of each scale to the overall correlation. The overall positive correlation is dominated by patterns at broad genomic scales.

Turning to the role of selection in shaping ancestry patterns, we next applied our wavelet methods to examine the spatial structure of the correlation between the proportion of inferred malinche-like haplotypes and the recombination rate along the genome. Recall that genetic drift will not produce any systematic associations between ancestry state and recombination rate, but that selection acting systematically against alleles from one ancestry will generate positive correlations between the recombination rate and the proportion of the ancestry being selected against. To match previously reported correlations, we analyze these correlations on the physical map of the genome. We find an overall positive correlation between malinche-like proportion and recombination that is largely driven by broad-scale patterns (Fig. 4b,c). While finer scales contribute progressively more to the overall correlation across the sampling interval (Fig. S9), this pattern appears to be driven by the increasing variance at finer scales rather than increases in the correlations at these scales (Figs. 4a, S10). Thus, while we observe strong broad-scale correlations consistent with selection acting against malinche alleles early on in the formation of the hybrid zone against, we do not find evidence that much of the change genome-wide ancestry patterns across this sampling interval (2006-2018) reflects selection against malinche alleles. Nonetheless, selection may still be shaping patterns of local ancestry; (Powell *et al*. (2021) reported evidence of contemporary selection against the malinche allele at a QTL for tail length in this population).

When viewed on the genetic map, we find that strong positive correlations are largely restricted to broad scales consistent with a recent origin of the hybrid population (Fig. S10b). We note however that we also detect weak but significant positive fine-scale correlations on the genetic map, e.g. at 2^−10^ Morgans, which would suggest selection acting on ancestry variation at finer scales than expected given the estimated age of the hybrid population. These finer scale correlations could possibly reflect older admixture present in the source populations that formed this hybrid population, or errors in either HMM similarity calls or recombination map at fine scales.

Assuming that any systematic relationship between recombination rate and ancestry proportion can only be generated by selection, we can use the percent of variation explained from a regression of ancestry wavelet coefficients on recombination rate wavelet coefficients to give an estimate of the percent of the variation in mean ancestry along the genome that can be attributed to selection at each genomic scale. Applying this logic, we find that ∼20% of variance at the broadest scales on both the physical and genetic map (i.e. >3.2 Mb, > 0.125 Morgans) can be attributed to selection. We consider this a lower bound estimate, given that it treats genomic similarity to *X. malinche* as a proxy for the locations of selected loci, and that it assumes a model where selection always acts against malinche alleles. This approach can easily be extended by including wavelet coefficients for other genomic features such as the density of coding base pairs as predictors in the regression. For simplicity and consistency across data sets we only show results using recombination rate as a predictor, though we note that, in this data set, including wavelet coefficients for coding sequence density as a predictor did not generally improve model fit.

### 3.4 Application to hybrid baboons in Amboseli

Genome-wide selection against hybrids has also been inferred in the case of hybrids between yellow baboons (*Papio cynocephalus*) and anubis baboons (*P. anubis*). Vilgalys *et al*. (2022) analyzed whole genome sequence data from 442 baboons sampled near the center of a hybrid zone in the Amboseli basin of Kenya. This population is comprised of a complex mixture of early and late generation hybrids (potentially reflecting recurrent admixture over hundreds of generations), with most individuals having majority yellow-like ancestry. Although overall levels of anubis-like ancestry have been gradually increasing in this region over time due to immigration of anubis-like individuals into the hybrid zone (Samuels and Altmann 1986; Tung *et al*. 2008; Vilgalys *et al*. 2022), Vilgalys *et al*. (2022) report positive genome-wide correlations between recombination rate and anubis-like ancestry, consistent with widespread selection against alleles carried on this ancestry background within the hybrid zone.

We took advantage of the complex history of hybridization in this system to further demonstrate how the power spectrum can reveal demographic trends in hybrid zones. We apply the wavelet variance decomposition to an interpolated estimate of the proportion of anubis-like haplotypes within diploids at a set of loci differentiated between allopatric reference panels of yellow and anubis baboons. Overall, the majority of spatial genomic variance is present at finer scales on the genetic map compared to the swordtail hybrid population, suggesting hybridization occurring over much deeper timescales (Fig. 5A). We also find that the power spectrum varies according to an individual’s genome-wide proportion of anubis-like ancestry, with the most anubis-like individuals showing greater ancestry variance at broad genomic scales. This pattern is consistent with the most anubis-like individuals in the sample being recent descendants of migrants with longer contiguous tracts of anubis-like ancestry, and implies multiple bouts of admixture rather than a single pulse. In sum, the power spectrum for ancestry state is consistent with this population being the product of both recent and historical hybridization events as previously inferred.

**Figure 5:**
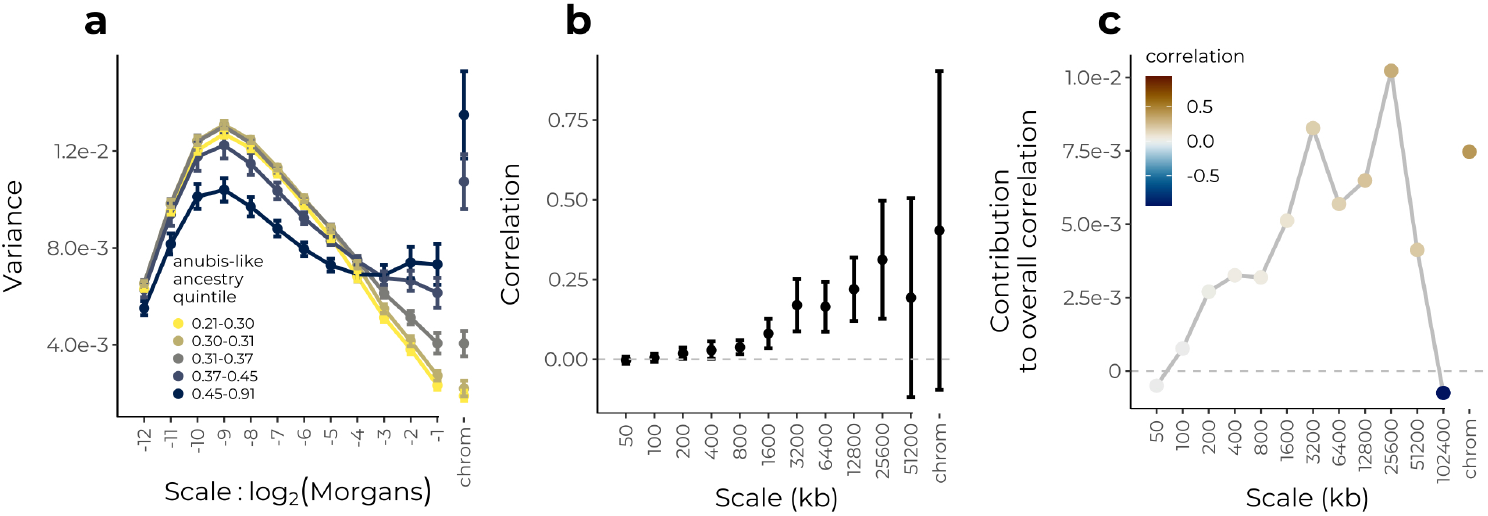
Wavelet analysis of hybrid genomes between yellow and anubis baboons in Amboseli, Kenya. **(a)** Power spectrum of the proportion of anubis-like haplotypes within diploids on the genetic map, stratified by quintile of genome-wide average anubis-like ancestry. Error bars are 95% confidence intervals using the standard error of the wavelet variance across individuals within each quintile. **(b)** Correlations at each spatial genomic scale (on the physical map) between wavelet coefficients for sample proportion of anubis-like haplotypes and wavelet coefficients for recombination rate. Squared values give an estimate of the proportion of variance in ancestry state explained by systematic selection against anubis-like alleles. Weighted-jacknife 95% confidence intervals shown for all but the largest scale which is present only on a single chromosome. **(c)** Contribution of each scale to the overall correlation. Although there is a strong negative correlation at scale 102400 kb, this scale is only present on chromosome 1 and does not contribute substantially to the overall positive correlation.

We next applied the wavelet methodology to examine the structure of a previously reported positive correlation between recombination rates and the proportion of anubis-like haplotypes along the genome (Vilgalys *et al*. 2022). We again find positive correlations across multiple scales on the physical map, with broad scales contributing the most to the overall correlation (Fig. 5B). Comparable to the swordtail population, we find that ∼20% of the variance in the proportion of anubis-like ancestry can be explained by systematic selection against anubis alleles, using just recombination rate as a predictor (Fig. 5B, squared values). When viewed on the genetic map, we find positive correlations across multiple scales consistent with ongoing selection against anubis-like ancestry, again with the strongest correlations at broadest scales (Fig. S11).

### 3.5 Application to Neanderthal ancestry in modern humans

As modern humans expanded out of Africa ∼ 60,000 years ago, they interbred with Neanderthals present in Eurasia (Green *et al*. 2010). Despite some Neanderthal variants having been adaptive (reviewed in Racimo *et al*. 2015), a number of studies have inferred that Neanderthal ancestry was on average deleterious in the human background, as Neanderthal-like haplotypes in modern humans are relatively depleted near conserved elements and in regions of low recombination (Sankararaman *et al*. 2014; Vernot and Akey 2014; Juric *et al*. 2016; Schumer *et al*. 2018).

We applied wavelet methods in order to understand the relative contributions of drift and selection in shaping variation in Neanderthal ancestry along the genome. Here we use three different call sets of Neanderthal-like haplotypes, inferred for the CEU 1000 genomes samples (Sankararaman *et al*. 2014; Steinrücken *et al*. 2018) and a large sample of modern Icelanders (Skov *et al*. 2020). Analyzing the power spectrum of the sample proportion of Neanderthal-like haplotypes on the autosomes for each set, we find reasonable agreement between our estimates and theoretical expectations under a model of a single pulse of neutral admixture 2,000 generations ago (Fig. 6a). Thus, while previously examined deserts of Neanderthal ancestry likely reflect early purifying selection against Neanderthal haplotypes, overall most of the variance in the frequency of Neanderthal-like haplotypes is on finer scales, largely consistent with the long term effects of genetic drift. While this is perhaps surprising given previously inferred negative fitness costs of introgression, we found in simulations of dispersed weak selection (assuming 10,000 loci contribute to a 20% fitness reduction in human-Neanderthal F1s, e.g. Harris and Nielsen 2016; Juric *et al*. 2016) generates only subtle deviations from the neutral power spectrum (Fig. S12). The sample average calls of different methods are in good agreement at broad scales (upwards of several Mb), but are only weakly correlated at finer scales of measurement (e.g. tens to hundreds of kb, Fig. 6b). This is expected given the low inferred proportion of Neanderthal ancestry and the old age of the admixture event. All of the methods have substantially reduced power to detect short fragments, particularly in the presence of recombination rate heterogeneity (Skov *et al*. 2018).

**Figure 6:**
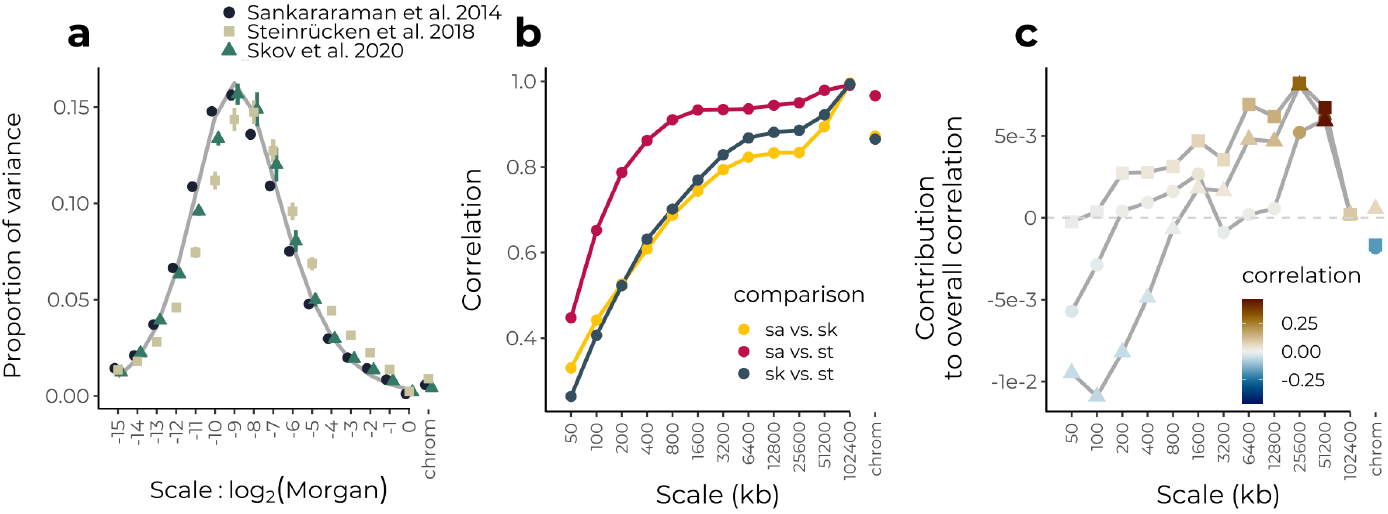
Wavelet analysis of inferred Neanderthal ancestry in modern humans. **(a)** Normalized power spectra of the proportion of Neanderthal-like tracts for three different studies (colored points) compared to a neutral expectation for 2,000 generations of admixture in a population size of 10,000 diploids (grey line). Error bars represent 95% confidence intervals from a weighted jackknife across chromosomes. **(b)** Correlations across scales between different similarity calls from different data sets. **(c)** Contribution of each scale to the overall observed correlation between the proportion of Neanderthal-like haplotypes and log-transformed recombination rates. Shapes correspond to different studies as indicated in panel (a).

Turning to correlations between the proportion of Neanderthal-like haplotypes and recombination rate, we find a consensus across data sets of positive correlations at broad scales on the physical map (Figs. 6c, S13). However, at the finest scales, we find significant negative correlations for some of the call sets. These are not due to a confounding correlation between recombination and gene density, which are negatively correlated at these scales (Fig. S14). The negative correlations could conceivably be generated by selection in late generations systematically favoring Neanderthal ancestry (as seen in our simulations of the case where introgressed variants are first deleterious but then later favored, Fig. 3D). However, we suggest they more likely reflect an inherent bias towards detecting Neanderthal-like haplotypes in regions of lower recombination. The estimated correlations at fine scales are variable across calling methods and appear at genomic scales where there are only weak correlations among sets of calls, suggesting these negative correlations are an artifact of the methods rather than underlying biological signal. Furthermore, we note that correlations at fine scales were highly sensitive to the posterior threshold applied in at least one data set, further suggesting that inference of ongoing or recent selection based on fine scale patterns of Neanderthal-like haplotypes is limited with current methods (Fig. S15).

## 4 Discussion

Wavelet analysis of admixed populations offers a useful look into the history of evolutionary forces acting in hybrid populations. We have shown with theory and simulations how a decomposition of the variance in ancestry state along the genome can be used to reveal the timing of genetic drift under a simple population genetic model of admixture. Further, we used simulations to illustrate how selection impacts the scale of both the variance in ancestry state and the correlation between introgressed ancestry and recombination rate. In total, these methods offer a compact summary of genome-wide admixture signals, and can inform a more general understanding of the role of selection in shaping patterns of introgression across the genome.

In applying the method to several systems, we can observe generalities in the genomic consequences of hybridization. Most notably, the observed positive correlations between introgressed ancestry and recombination rate are largely dominated by patterns at broad genomic scales. Simulations indicate that these patterns establish rapidly in the earliest generations after hybridization when selection acts against multiple alleles from one ancestry across the genome. Thus, this evidence accords with our understanding that selection is strongest on early generation hybrids (Barton 1983; Harris and Nielsen 2016; Veller *et al*. 2023), and suggests that this effect can be detected even hundreds or thousands of generations after an admixture event. In all three cases, a reasonable proportion of the variation in ancestry state along the genome at the broadest scales is attributable to recombination rate variation (roughly 10-20%). As these estimates are a lower bound on the contribution of selection, we can say that selection plays a key role in shaping the genomic composition of early generation hybrid populations, with lasting effects. Weaker correlations at finer genomic scales also agree with our understanding that the strength of selection on hybrids dramatically decreases over generations (Veller *et al*. 2023), and suggests that genetic drift may be the dominant force shaping fine-scale genomic ancestry patterns. A related approach to estimating the contribution of selection to ancestry patterns across scales would be to apply these analyses to the correlation in ancestry state between independent replicates of hybrid populations derived from the same source populations. This approach may be more powerful in that locations and effects of selected alleles are internally matched when similar parental sources repeatedly hybridize under similar ecological conditions.

In demonstrating the effects of selection on wavelet-based statistics, we considered a simple additive model of selection against alleles at multiple loci carried in one ancestry background. However, the wavelet methods themselves are agnostic to the form of selection. Alternative models of selection that incorporate models of dominance and epistasis could generate different signatures in the wavelet statistics, such that these methods could potentially be applied towards distinguishing among models. For instance, Harris and Nielsen (2016) found that if deleterious mutations in human and Neanderthal haplotypes are largely partially recessive, the direction of selection on introgressed Neanderthal ancestry in human populations could change through time due to the contrasting effects of purifying selection and selection for heterosis (see also Kim *et al*. 2018). We have shown here that such a reversal in the direction of selection on one ancestry can in principle be detected using the wavelet decompositions (Fig. 3d). One parameter explored here is the total number of loci under selection in hybrids. Using the same additive model with the total additive strength of selection held constant, we found that the wavelet decompositions vary with the overall number - and correspondingly the density - of sites under selection in hybrids. Importantly, a high density of sites under selection is required to generate fine-scale correlations between introgression and recombination. Thus, the scale at which correlations between introgressed ancestry and recombination rate are generated is indicative of the scale at which selected loci are distributed in the genome, and thus their total number.

An important caveat to these methods is that any systematic biases in the input data, including estimation of recombination rates and local ancestry inference, will be propagated into the wavelet transform. For example, biased detection of introgressed fragments toward low recombination regions may generate spurious signals of selection favoring introgressed ancestry genome-wide. We have suggested this may be the case for Neanderthal introgression into humans, where we measure negative correlations over fine genomic scales (Fig. 6C). These potential biases, together with inconsistent patterns across sets of calls and high sensitivity to posterior probability thresholds also suggest more generally that inferences of selection on Neanderthal ancestry relying on fine-scale genomic patterns might need to be revisited (Juric *et al*. 2016; Telis *et al*. 2020). Other issues such as the need for phased reference panel haplotypes and the potential for model mis-specification limit wider applicability of local ancestry inference with HMM-based methods in non-model systems.

One promising direction to overcoming these limitations is that wavelet transform can be applied to single-locus admixture statistics (Pugach *et al*. 2011; Sanderson *et al*. 2015) to avoid directly inferring the boundaries of ancestry tracts. While these statistics contain additional noise unrelated to the admixture process, future work could develop a theoretical framework for applying these methods to single-locus admixture statistics, thereby avoiding biases arising from local ancestry inference with HMM methods. Additional theory is also needed to better understand the impact of selection on the wavelet variance and correlation decompositions, and to extend these ideas to scenarios with ongoing gene flow and hybrid zones.

We now appreciate that introgression is a common feature of eukaryotic genomes, and the proliferation of genomic sequence data presents an opportunity to study hybridization events even in the ancient past. Combined with the increasing availability of more complete genome assemblies and recombination maps, wavelet approaches should enable patterns in the strength and time scale of selection on hybrids to emerge across systems.

## 5 Materials and Methods

### 5.1 Simulation and data processing

All simulations were performed in SLiM 4.0 (Haller and Messer 2019). Each locus in our simulations represented a genomic window of fixed physical length (e.g. 50kb). Recombination rates between adjacent windows were modelled off a genetic map of the human autosomes estimated in Halldorsson *et al*. (2019). In simulations with selection, we fixed deleterious alleles in one source population at 10,000 loci placed uniformly at random on the physical map in each replicate run. We assumed a toy model of polygenic selection against introgressed ancestry, where the fitness of individual *i* (*w*_*i*_) declines linearly with the fraction of introgressed alleles that the individual carries (*p*, with the slope given by *S*: *w*_*i*_ = 1 − *pS*. Similar models were studied in Barton (1983), Barton and Bengtsson (1986), and Veller *et al*. (2023). For the analyses shown in the main text we set *S* = 1, corresponding to F1 hybrids having a relative fitness of 0.5. Simulations used tree sequence recording (Haller *et al*. 2019), and ancestry along the genome was extracted from the tree sequences using *tskit* (https://tskit.dev/software/tskit.html).

For downstream wavelet analysis, we require signal values at evenly-spaced positions, either on a physical or genetic map. For our simulations where simulated loci represented physical genomic windows of fixed length, we thus performed interpolation of ancestry and recombination signals to a grid of evenly-spaced positions on the genetic map.

### 5.2 Wavelet analysis

For all wavelet analyses, we used the Maximal Overlap Discrete Wavelet Transform with Haar wavelets (Percival and Walden 2000) implemented in the R package *waveslim* (Whitcher 2022). Further background on the wavelet methods are provided in the appendix.

As the wavelet transform only operates on contiguous signals, we perform the variance and correlation decompositions to each chromosome separately and then combine results across chromosomes in one of two ways. Due to heterogeneity in chromosome length, not all scales will be present on all chromosomes. Thus, for estimating raw magnitudes of the wavelet variance, we average over only those chromosomes for which a given scale is present, taking a weighted average where the weight is chromosome length. Separately, we calculate the proportion of total genomic variance contributed each scale by assigning a variance value of zero to those chromosomes for which a given scale is not present, and then performing the same weighted average across chromosomes. Values are then adjusted to account for the proportion of total genomic variance due to variance among chromosomes in their mean values of a signal. The among-chromosome portion is also a weighted average weighted by chromosome length. Note that the calculations shown in this manuscript remove so-called boundary coefficients (further details in appendix), which leads to unbiased estimates of wavelet variance, but means that the proportion of total genomic variance at each scale given is approximate.

Generic functions to perform all the analyses in this paper are contained in the R package *gnomwav* available at https://github.com/jgroh/gnomwav. The function *gnom var decomp* returns the genome-wide variance decomposition for any signal, including both forms of averaging over chromosomes. Likewise, *gnom cor decomp* returns genome-wide wavelet correlations for each scale. The contribution of each scale to the overall correlation between signals can be obtained from the output of these two functions. These functions are not specific to the investigation of admixture and should be broadly applicable to many genomic signals of interest.

### 5.3 Swordtail analysis

We used genomic similarity calls from Powell *et al*. (2021), consisting of posterior probabilities for diploid genotypes matching reference panels of *X. malinche* or *X. birchmanni* at a set of SNPs that are highly differentiated between the species. The frequency of the minor parent allele (*X. malinche*) for each individual was taken as a weighted average of the posterior probabilities of being homozygous and heterozygous for matching *X. malinche*, i.e. 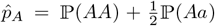. We interpolated these estimates to a grid of 2^−12^ Morgans, chosen such that the density of SNPs was on average greater than the density of interpolated grid points. For analyses on the physical map, we interpolated ancestry estimates to a 50kb grid.

We obtained positions of SNPs at which genomic similarity to reference populations was called (ancestry SNPs) along an LD-based recombination map (Schumer lab, pers. comm.), using the median values of 2*Nr* per SNP interval to get cumulative genetic distances along each chromosome. Since these estimates are given in units of the population-scaled recombination rate, i.e. *ρ* = 2*N*_*e*_*r*, we converted distances to Morgans using an estimate of 2*N*_*e*_ from the slope of a regression between the genetic lengths of chromosomes estimated from a crossover map and those estimated from the LD map. As we observed extreme outliers in values of *ρ*, we truncated the distribution of 2*N*_*e*_ at 0.005, corresponding to 1.6% of the total genome (a similar threshold was also applied in Schumer *et al*. 2018). We applied the lowest threshold possible beyond which we saw relatively stable estimates of *N*_*e*_, and we also observed a significant improvement in the fit of the above regression using the chosen value. We also found that results applying this threshold show better agreement with a previously inferred age of the hybrid zone (Schumer lab, pers. comm.).

### 5.4 Amboseli baboon analysis

We used genomic similarity calls from Vilgalys *et al*. (2022), i.e. posterior probabilities for diploid genotypes matching reference panels of *P. anubis* or *P. cynocephalus* at a set of SNPs that are differentiated between the species. Estimates were interpolated as described above using an LD-based recombination map. As we again observed extreme outliers in values of *ρ*, we applied a threshold at 0.01 (corresponding to 1.4% of the total genome). To visualize the wavelet variance spectrum on a genetic map, we converted genetic lengths to units of Morgans as described above, using genetic lengths of chromosomes from Rogers *et al*. (2000).

### 5.5 Neanderthal introgression analysis

We compared results using three separate estimates of Neanderthal-like haplotype frequency, from Skov *et al*. (2020), Sankararaman *et al*. (2014), and Steinrücken *et al*. (2018). Data for the former were obtained in the supplementary data associated with the article, and data for the latter two were downloaded from https://web.cs.ucla.edu/~sriram/software-data.html and https://dical-admix.sourceforge.net/, respectively.

We interpolated the estimates to 50kb windows on a physical map of the genome, 2^−12^ Morgans on the genetic map. For the frequency estimates from Skov *et al*. (2020), we used the sum across identified archaic fragments of a weighted average of the frequency of each fragment in the sample of Icelanders, with the weight being the portion of the window covered by the fragment. For the data from Sankararaman *et al*. (2014) and Steinrücken *et al*. (2018), we used Neanderthal allele frequency estimates in the CEU sample of the 1000 genomes project (2N=170). For these two studies, we tried two separate measures of Neanderthal-like haplotype frequency. First, we directly used marginal posterior probabilities of a site matching Neanderthal, averaged across individuals in the sample, e.g. column 11 in the output files provided by Sankararaman *et al*. (2014). Next, following the analyses of the original authors, we applied a threshold to the marginal posterior probabilities, calling sites with marginal posterior probability above 0.90 from Sankararaman *et al*. (2014) and above 0.42 from Steinrücken *et al*. (2018) as Neanderthal-like, then taking the average of calls across haplotypes (e.g. column 15 in the files provided by Sankararaman *et al*. (2014)). We also tried the analyses in coordinates of both hg19 and hg38, using liftOver to convert estimates across assemblies. For analyses using hg38, we used the recombination map from Halldorsson *et al*. (2019). For analyses using hg19, we used the recombination map provided directly in the available output files from Sankararaman *et al*. (2014). Results shown in the manuscript are in hg38 coordinates.

We note that recombination rates in humans are likely estimated with greater resolution relative to the data sets above; consistent with this we found considerably greater variance in the recombination rate across scales, and thus used log-transformed recombination rates for the analyses shown in this paper.

## 6. Acknowledgements

We are grateful to Carl Veller, Michael Turelli, and the Schumer lab for comments on the manuscript, and to Rasmus Nielsen, Molly Schumer, and members of the Coop lab for helpful discussions. Funding was provided by the National Institutes of Health (NIH R35 GM136290 awarded to GC) and the National Science Foundation Graduate Research Fellowship (NSF 1650042 awarded to JG).

## Supplementary Material

### S1

As wavelets have not been widely used in population genetics, here we give some basic background on the Discrete Wavelet Transform. See Percival and Walden (2000) for a comprehensive treatment on theory and practical applications of the wavelet transform.

### S1.1 Discrete Wavelet Transform (DWT)

The starting point for the DWT is a signal, i.e. a set of measurements that are ordered in space or time. In our case, these are measurements of ancestry state taken along a contiguous stretch of DNA, e.g. a single chromosome. We will write this ancestry signal as the vector **x** with entries *x*(𝓁),𝓁 = 0, 1, 2, …, *L* − 1, where *L* is the total length of the chromosome being analyzed. The values of 𝓁 must be evenly spaced in units defined by the resolution of measurement. For example, if ancestry state is measured at every nucleotide position, *L* would equal the number of nucleotide positions in the chromosome. If the ancestry state is instead measured in windows of 10kb along a chromosome, *L* would equal the number of such windows that fit on the chromosome. We will impose an additional restriction that *L* is a power of two (this will be relaxed later).

The DWT is an orthogonal transformation, where we project the ancestry signal onto a new set of coordinate axes. Instead of viewing the value of the ancestry signal at positions 𝓁, we will instead view the data on a new set of coordinate axes where axes have the interpretation of measuring the magnitude of *change* in the ancestry signal at a particular genomic scale (defined in more detail below), and at a particular location. The DWT can be represented as a linear transformation as follows:

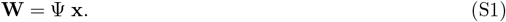

Above, **x** is a column vector, multiplied on the left by a square matrix Ψ to yield a new column vector **W**, whose entries are the wavelet coefficients. The rows of Ψ are thus the basis vectors of the new coordinate system. The first *L* − 1 of these are known as *wavelets*, and the *L*^th^ is known as the scaling filter. These are defined below. In practice, the DWT is accomplished using a fast algorithm known as the pyramid algorithm (Percival and Walden 2000).

#### S1.1.1 Haar wavelets

Each wavelet *ψ*(𝓁) is written as a function of the genomic position 𝓁, and so is defined on the same set of values as *x*(𝓁). When the values of *ψ*(𝓁) are plotted against the values of 𝓁, the function resembles a wave in the sense that the values grow and decay. However the wave is ‘small’ in the sense that this oscillatory behaviour may be limited to a subset of the domain. For example, a sine wave oscillates infinitely over its domain and so does not obey this second property. This notion is formalized in the following two properties:

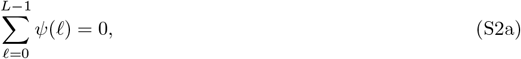

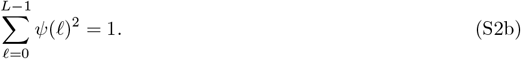

Any function with these properties could be used as a wavelet. However, for a given DWT analysis, all of the wavelets in the matrix Ψ belong the same family. The wavelets of the same family are all related through a set of shifting and scaling operations. From here on we specifically focus on Haar wavelets. A Haar wavelet is a piecewise constant function that is non-zero over a finite portion of the sequence, and has a single transition from positive to negative values (see Fig. 1 for examples). Each Haar wavelet is indexed by two parameters, *j* and *k*. (Note in the main text we index wavelets using *λ* and *i* in place of *j* and *k* for ease of description, but the two notations are equivalent). The parameter *j*, or the level of the wavelet, corresponds with the length of sequence over which the Haar wavelet is non-zero, and the parameter *k*, or shift, corresponds with where in the sequence the non-zero portion is located. Formally, the set of Haar wavelets used in the DWT is defined as follows:

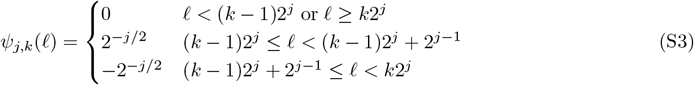

where *j* = 1, 2, … log_2_ *L* and *k* = 1, …, *L/*2^*j*^. Haar wavelets of level *j* only take non-zero values in one interval of the domain, which is of length 2^*λ*^. We define the *scale* of a wavelet, *λ* to be equal to half of this interval, i.e. *λ* = 2^*j*−1^, so we can say the wavelet changes from positive to negative over a scale of *λ* (this definition corresponds with the description of *λ* in the main text). Thus, the set of scales that the DWT yields information about are a dyadic series, that is, successive scales differ by a factor of two. We also see that the number of wavelets at a given level, *L/*(*λ* + 1), is simply the number of wavelets at that level that fit onto the chromosome with no overlap in where they take non-zero values. Because of our restriction that *L* must be a power of two, this is always whole number. Thus, the number of wavelets decreases by two for successively larger scales. If we order the wavelets at a given level with their non-zero portions arranged adjacent from left to right, *ψ*_*j,k*_ is the *k*^*th*^ wavelet in this ordered set. In this sense, the parameter *k* describes where in the domain the non-zero portion of the wavelet is located.

The definition given above satisfies a third property of the wavelets used in the DWT, which is that they form an orthogonal set. Formally,

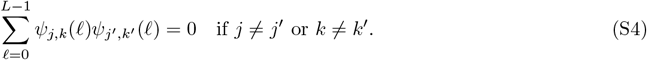

As a toy example, for a very small chromosome with *L* = 8, the wavelets used in the DWT are:

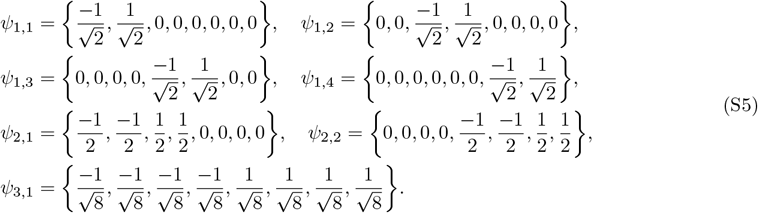

There is additionally one *scaling filter, G*(𝓁), which is constant valued, 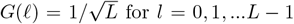. Because of this, *G* does not obey property S2a, but it does obey properties S2 and S4.

#### S1.1.2 Wavelet coefficients

We have now seen that the rows of Ψ are the wavelets *ψ*_*λ,k*_(𝓁), along with the scaling filter *G*(𝓁) as described above. The *n*^*th*^ entry of **W** is an inner product between the *n*^*th*^ row of Ψ and the vector **x**. The inner product between a wavelet and **x** yields a wavelet coefficient. Because the mean of any wavelet is zero, a particular wavelet coefficient is proportional to the covariance between the ancestry signal and the associated wavelet:

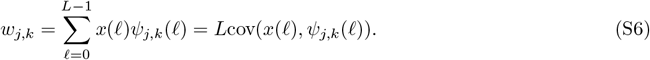

The inner product of the scaling filter and **x** produces the *scaling coefficient* which is proportional to the mean of *x*(𝓁):

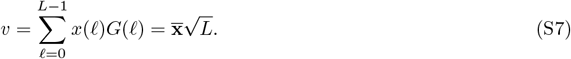

If we sort the rows Ψ by increasing *j* and then *k*, and finally the scaling filter *G*, we can expand the matrix multiplication of Eqn. S1 using Haar wavelets:

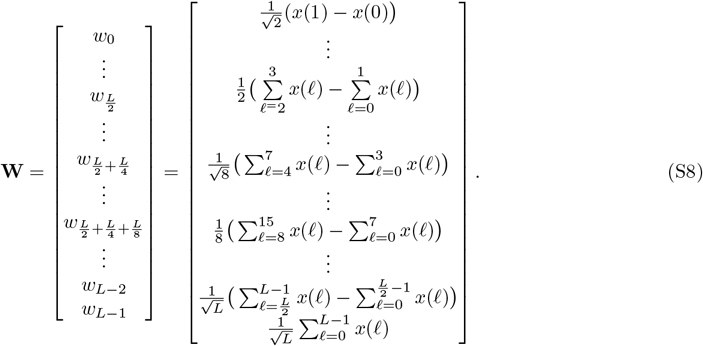

From this we can see that the magnitude of a particular wavelet coefficient produced with a Haar wavelet is proportional to the magnitude of change occurring between two adjacent windowed averages of the ancestry signal, where the window length is the scale of the associated wavelet.

Thus, we are now viewing **x** in a new set of coordinates that describe (i) fluctuations around that mean value of **x** and different scales and locations, and (ii) the mean value of **x** itself.

#### S1.1.3 DWT wavelet variance decomposition

We can use the wavelet coefficients to understand what portion of the total variance across loci in ancestry state can be attributed to variation at different genomic scales. Since the DWT is an orthonormal transformation, we have the following property: ∑_𝓁_*x*(𝓁) = ∑_𝓁_*W*(𝓁)^2^. Using this property and the property that 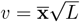, we can write the sample variance of **x** as a sum of contributions from different scales, indexed by *λ*:

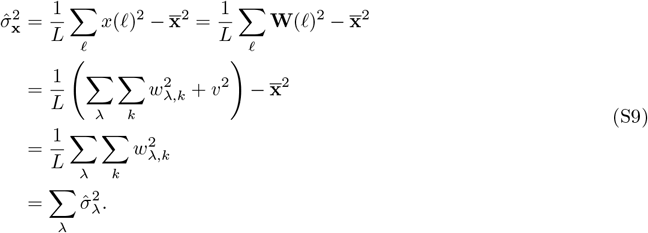

#### S1.1.4 Wavelet covariance decomposition

Analogous to the wavelet variance, we can compute the wavelet covariance at scale *λ* for two signals *x* and *y*:

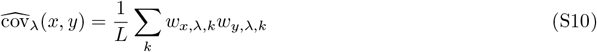

and for the DWT the sum of these gives an exact decomposition of the total covariance:

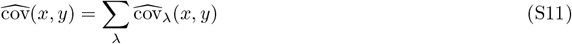

### S1.2 Maximal Overlap Discrete Wavelet Transform (MODWT)

The DWT is restricted to signals that are powers of two, and the coefficients produced by the DWT are not shift-invariant. In other words, the set of coefficients resulting from performing the DWT on a shifted signal (moving every value by the same amount in one direction, and treating the signal as circular) are not the same set of coefficients resulting from the DWT on the original signal.

The MODWT is a related transform that is both well-defined for signals of any length and shift invariant (Percival and Walden 2000). In essence, it results from circularly shifting **x** and performing the DWT, resulting in a vector of wavelet coefficients **W**_*λ*_ of length *L* for each *λ*, representing localized changes in adjacent averages taken in windows of size *λ*, as well as one vector of scaling coefficients **V** of length *L* corresponding to average values of the signal in windows of size ⌊log_2_ *L*⌋. One consequence of this procedure is that the first *λ* elements of **W**_*λ*_ and **V** are so-called *boundary coefficients* that describe changes between discontiguous portions at the beginning and end of the signal. Different approaches for dealing with these coefficients when calculating wavelet variances are available (described in Percival and Walden). In our application, we have omitted boundary coefficients in our calculation of wavelet variances, but including boundary coefficients produced qualitatively similar results.

Also in contrast to the DWT, the MODWT is not an orthogonal transformation, as there is redundancy among neighboring wavelet coefficients within each level. In other words, wavelet coefficients will often be auto-correlated along the signal. This does not however bias the estimates of the wavelet variances, and we use MODWT as it produces less noisy estimates of wavelet variances. Similar to the DWT, the MODWT gives a complete decomposition of the sample variance as follows:

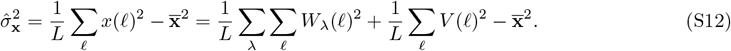

We define the portion 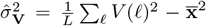 as the scaling variance, and note that unlike the wavelet variances, it’s calculation requires subtracting the squared mean of coefficients. The scaling variance has a different interpretation than the wavelet variances as it is associated with variance of average values of the signal taken in windows of size ⌊log_2_ *L*⌋, rather than being a variance associated with changes. Similarly, in computing the covariance decomposition shown above for the DWT, the exact decomposition includes the covariance of scaling coefficients for *x* and *y*. In contrast to the wavelet covariances, the scaling covariance subtracts the product of the means of the scaling coefficients of the two signals.

## S2

In this section we derive expectations of the wavelet variance for ancestry state along the genome under a neutral single-pulse model of hybridization.

### S2.1 Expectation for wavelet variance of sample mean ancestry

We write the mean ancestry at position 𝓁 in a sample of size *M* chromosomes as follows

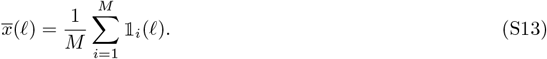

where 𝟙_*i*_(𝓁) is an indicator random variable; 𝟙_*i*_(𝓁) = 1 if haplotype *i* carries an introgressed allele at locus 𝓁 and is zero otherwise. We will later refer to the ‘recipient’ and ‘donor’ populations as populations 0 and 1, respectively, but the choice is arbitrary with respect to the admixture proportions.

From Eqn. S9, the expected contribution to the total variance of mean ancestry 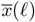 along the sequence associated with level *λ* is given by

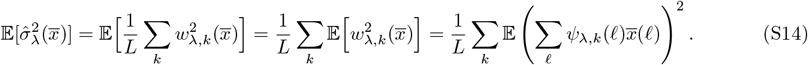

We can approximate the sum over the *L* loci as a continuous integral over a region of length *L*. Then,

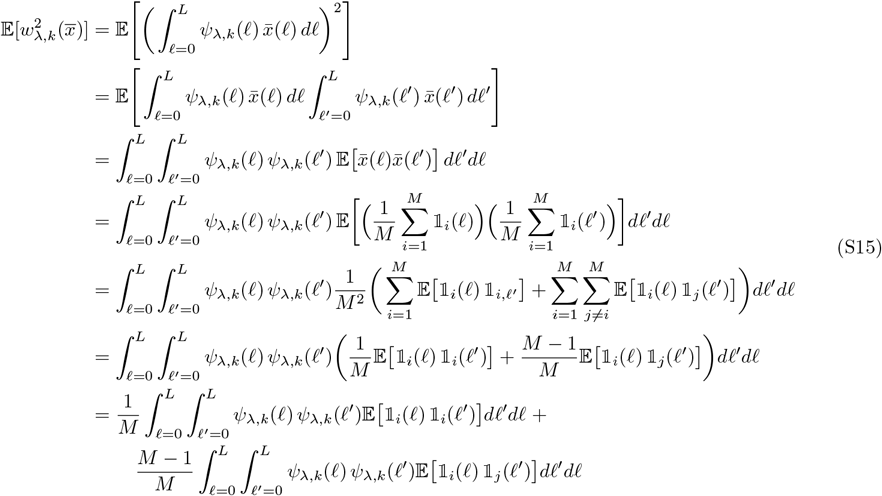

We can interpret 𝔼[𝟙_*i*_(𝓁) 𝟙_*j*_ (𝓁′)] as the probability that haplotype *i* inherits from the donor population at locus 𝓁 and individual *j* inherits from the donor population at locus 𝓁′. From (S15), we also see that for a general statistic, the wavelet variance of the sample mean is equal to a weighted average: 1*/M* times the average wavelet variance for single chromosomes, plus (*M* − 1)*/M* times the average wavelet covariance of pairs of chromosomes, where *M* is the sample size.

### S2.2 Neutral case

Under neutrality, both of the expectations in (S15) will only depend on the distance between 𝓁 and 𝓁′, and not their exact locations, so there is no dependence on *k*. Thus, we have

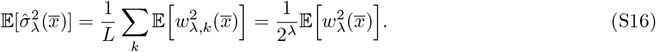

The two expectations in (S15) can then be calculated under a coalescent model using a two state continuous time Markov chain that runs backward in time from the present. Let state *X* = 1 represent the alleles at loci 𝓁 and 𝓁′ being present on the same haplotype, and state *X* = 2 represents the two alleles being present on separate haplotypes. The transition rate matrix is given by the coalescent approximation with time scaled to units of 2*N* generations (*τ* = 2*Nt*),

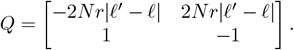

The transition probability matrix is given by

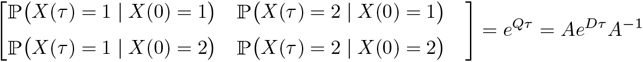

with *D* containing the eigenvalues of Q on the diagonal and the columns of *A* containing the corresponding right eigenvectors so that

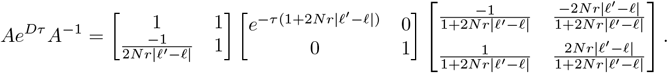

For an initial admixture proportion of *α*, two alleles present on the same haplotype at the time of admixture come from the donor population with probability *α*, while two alleles present on different haplotypes at the time of admixture come from the donor population with probability *α*^2^. We then have

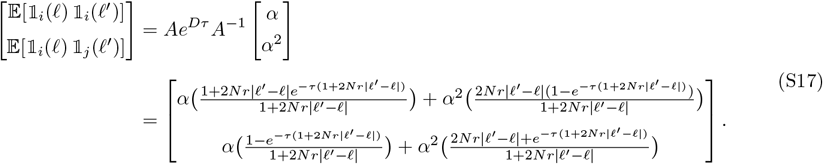

Note in the above that if we set *t* = 0 we get 𝔼[𝟙_*i*_(𝓁) 𝟙_*i*_(𝓁′)] = *α* and 𝔼[𝟙_*i*_(𝓁) 𝟙_*j*_ (𝓁′)] = *α*^2^ as expected. In this case, since *ψ*_*λ,k*_(𝓁) integrates to zero, the integrand in (S15) becomes an even function with no dependence on the distance between loci and we get 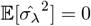. This makes intuitive sense; with no time for recombination after the initial admixture, there is zero variance at all spatial scales as introgressed alleles are in complete linkage. In the limit as *N* → ∞, both expectations depend only on the recombination process and we have

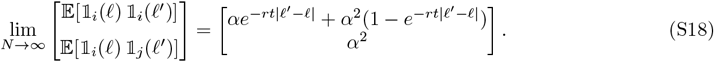

In this particular case the wavelet variance of the mean is equivalent to the wavelet variance for a single haplotype, since drift cannot induce any covariance among haplotypes.

These expectations in general can also be found under an any demographic model where the population change size changes through time as a piecewise constant function of time. The length of the total time interval between the present and the time of admixture, *τ*, is subdivided into *n* component internals of lengths *τ*_1_, *τ*_2_, …*τ*_*n*_ which sum to *τ*. The probability transition matrix over the entire interval is the result of multiplying probability transition matrices for subsequent subintervals on the right (moving backward in time):

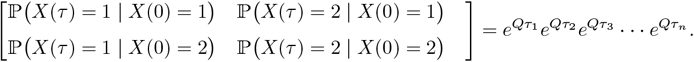

### S2.3 Discrepancy for early generations

In early generations after admixture, the coalescent provides a poor model for the distribution of ancestry tract lengths, because the fixed pedigree constrains which ancestors can be inherited from (Liang and Nielsen 2014). We thus expect error in the calculations above for the first few generations after hybridization. To illustrate this, we calculate the expected wavelet variance for the F2 generation, assuming no selection or assortative mating in the F1 generation. To match our Wright-Fisher simulation framework, we imagine that two populations mix with proportions *a* and 1 − *α*. In the F1 generation, individuals are either descended from a mating between two individuals from the same source population (events A, B with probabilities *α*^2^ and (1 − *α*)^2^, respectively), or they are F1s (event C) with probability 2*α*(1 − *α*). We then have

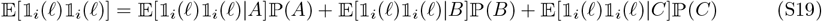

Using Haldane’s map function for the probability that a chromosome produced through meiosis in an F1 is not recombinant at 𝓁 and 𝓁′, this becomes

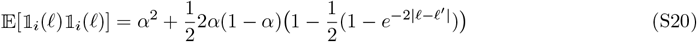

For 𝔼[𝟙_*i*_(𝓁) 𝟙_*j*_ (𝓁)], we only need to consider whether the lineages coalesce in the previous generation:

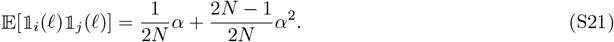

Using the expectations in Eqns. S20 and S21 in the integrand for the wavelet variance calculation, we find that this provides a much better fit for the simulated wavelet variance in the F2 generation compared to the coalescent model specified by Eqn. S17 (Fig. S1).

**Figure S1:**
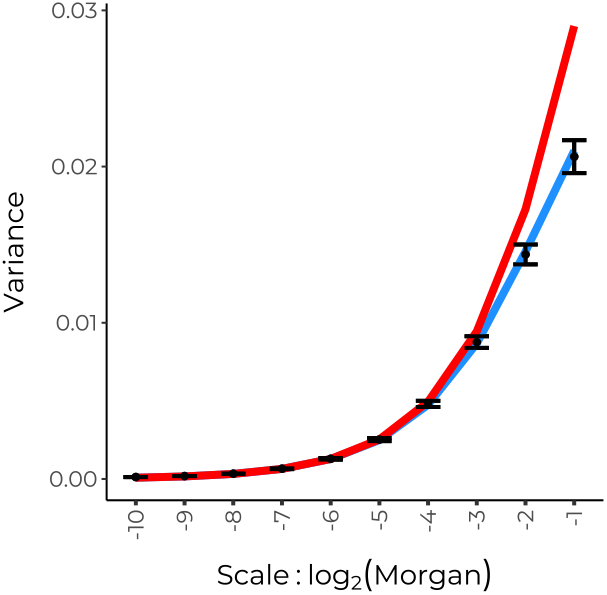
Black points and error bars show mean wavelet variances and 95% confidence intervals for ancestry state along single chromosomes in the F2 generation from 100 replicate simulations of a 50/50 mixture between two populations. Red line shows expected wavelet variance using the coalescent model. Blue line shows expected wavelet variance calculated for the F2 generation by conditioning on the pedigree.

**Figure S2:**
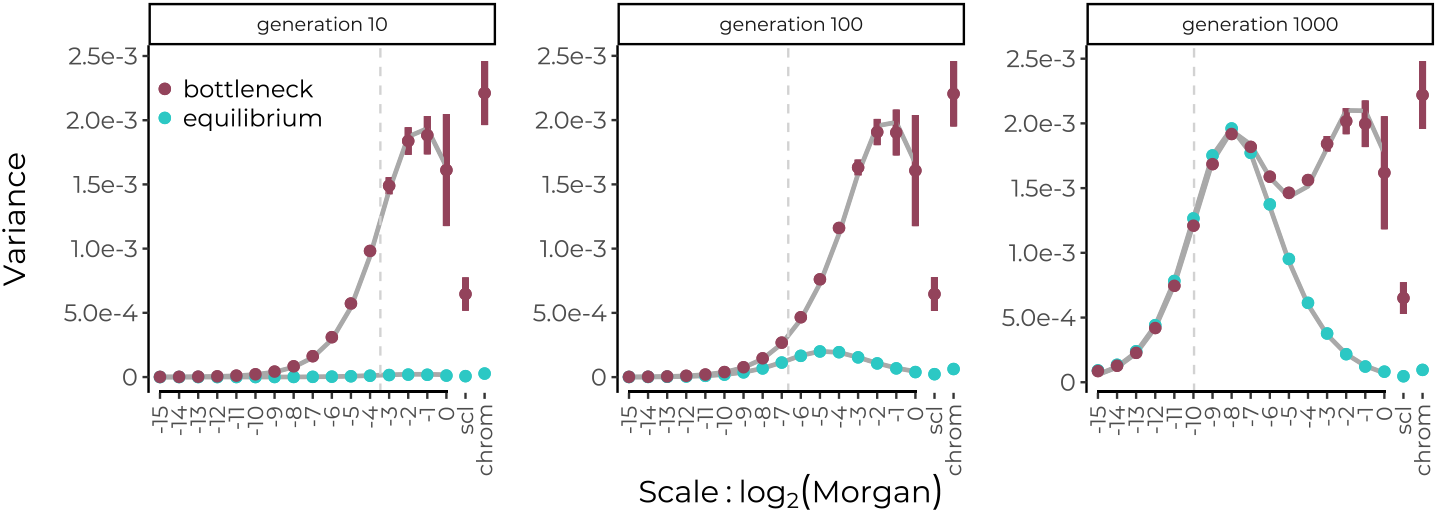
Simulated power spectrum of ancestry proportion in a hybrid population undergoing genetic drift compared to theoretical expectations. We simulated (using SliM, Haller and Messer, 2019) a population of constant size 2N=20000 (light blue) and a population that undergoes a bottleneck to 2N=200 for just the first 10 generations of recombination in hybrids, then expands to 2N=20000 (maroon). Points and error bars show means and 95% confidence intervals across 20 replicate simulations. Solid grey lines show theoretical expectations. Also shown are scaling variances collapsed into a single category ‘scl’ and chromosome-level variance. Vertical dotted lines are placed to indicate the expected distance between recombination breakpoints that have accrued along a single chromosome since the hybridization pulse. In contrast to the figures shown in the main text, simulated loci are evenly spaced on a genetic map of the human autosomes, thus no interpolation is required and the estimates closely match the theoretical expectation.

**Figure S3:**
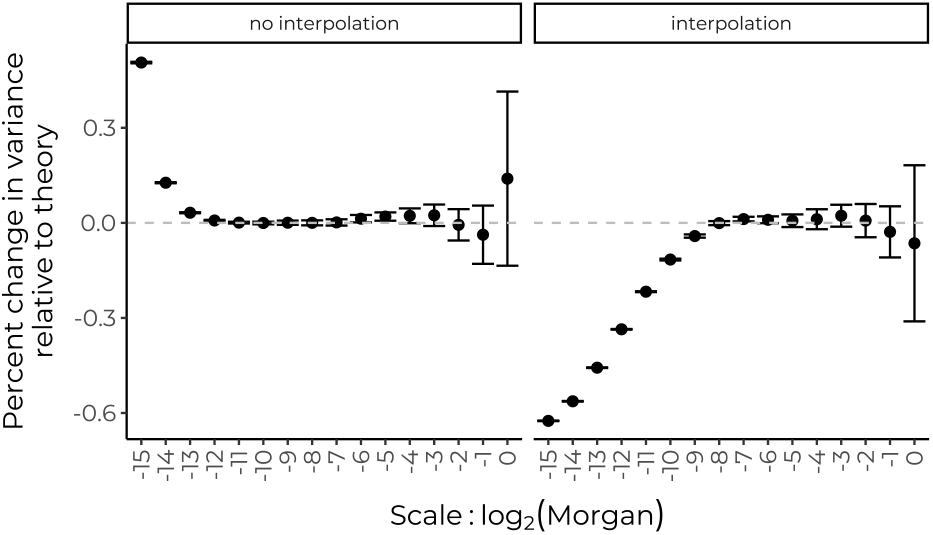
The effect of ancestry state interpolation on the power spectrum. Because informative markers are generally not evenly spaced, to perform the wavelet transform we must interpolate ancestry to evenly-spaced positions along the chromosome, either on a physical or genetic map. To see the scale of noise generated by interpolation, we compare simulated power spectra vs. theoretical expectations (Appendix B) for two cases after 1000 generations of mixture. **(Left)** Simulated loci are evenly spaced on the genetic map, thus no interpolation is required. In this case, deviations from the neutral expectation occur only at the finest scales, possibly reflecting numerical error. **(Right)** Simulated loci occur every 50kb on a physical map of the human autosomes, and ancestry is then interpolated to a grid on the genetic map to perform the wavelet transform. In this case, interpolation causes a downward bias over the left half of the power spectrum.

**Figure S4:**
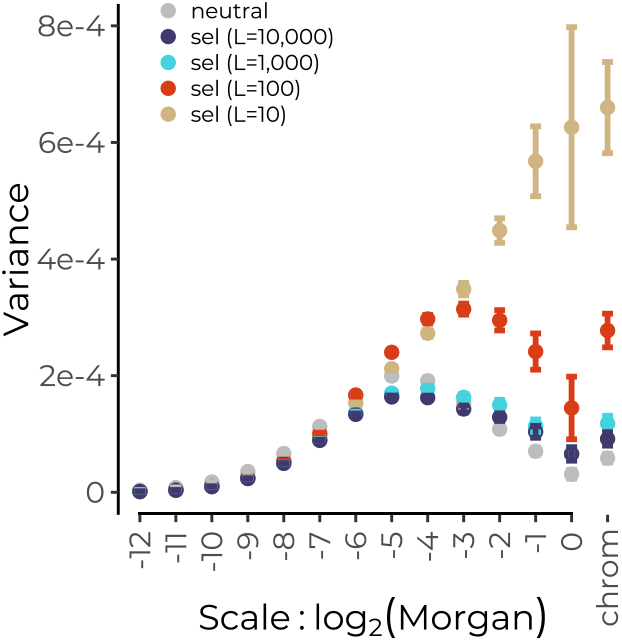
Selection against multiple alleles carried in one ancestry background distorts the wavelet variance decomposition of introgressed ancestry relative to the neutral expectation. We ran simulations of a 50/50 population mixture with selection acting against alleles from one ancestry at varying numbers of loci (10, 100, 1000, and 10000). In each case, the total strength of selection against F1 hybrids is held constant, where F1s have a 50% fitness reduction relative to the local source population. With fewer numbers of loci, there is stronger per locus selection concentrated in fewer regions, and selection at these loci generates increased broad-scale variation among regions harboring the deleterious loci and regions with mostly neutral variation.

**Figure S5:**
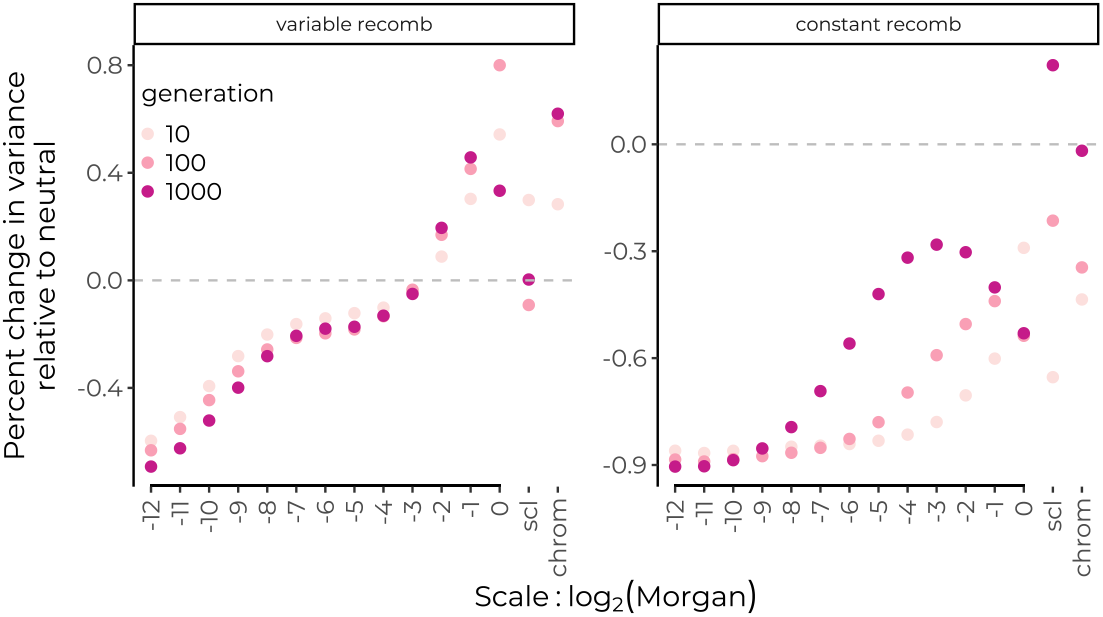
Genome-wide selection against alleles from one ancestry distorts the power spectrum. **(Left)** Selection on a heterogeneous recombination landscape modelled off the human autosomes generates greater variance at broad scales and lower variance at fine scales relative to the neutral expectation. Selection acts additively against alleles from one ancestry at 10,000 loci placed uniformly at random on the physical map, each selected allele has a selection coefficient of *s* = 5*e* − 5. The power spectrum of ancestry is compared to that from neutral simulations that use the same interpolation procedure. Thus, the distorion here is due to selection per se and not interpolation. The reduction in variance at fine scales after only 10 generations can be explained by the fact that selection brings the average introgressed ancestry proportion closer to zero. **(Right)** Here, simulated loci are placed evenly on a genetic map and results are compared to neutral simulations that do the same. Thus, although the overall variance is reduced, we see that selection distorts the power spectrum in a similar manner even in the absence of recombination rate heterogeneity.

**Figure S6:**
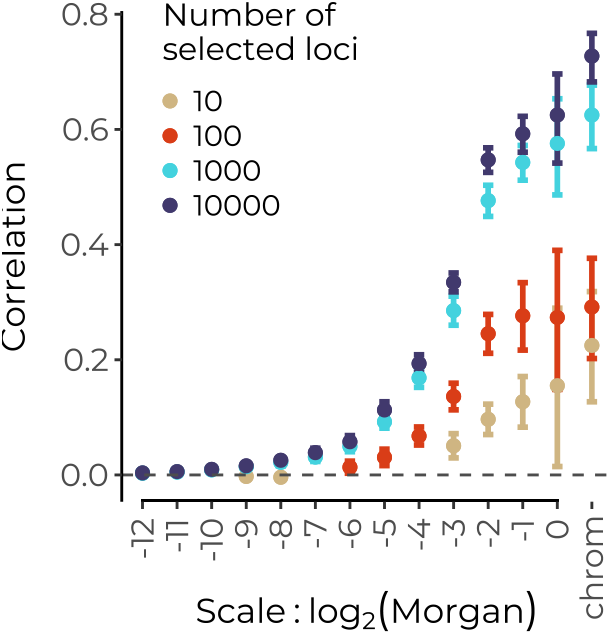
Selection against introgressed alleles generates correlations with recombination at varying scales according to the total number, and thus the spacing between selected loci. In simulations where the total strength of selection is held constant but distributed over varying numbers of loci, we find that in generation 1000, correlations with recombination are present at varying scales. With 10,000 loci under selection, fine scale recombination rate variation generates fine scale correlations between introgressed ancestry and recombination. With only 10 loci under selection, fine scale variation in recombination rate does not influence the rate of purging of deleterious introgressed alleles, so correlations at finer scales do not establish. Error bars represent 95% confidence intervals across 20 replicate simulations runs.

**Figure S7:**
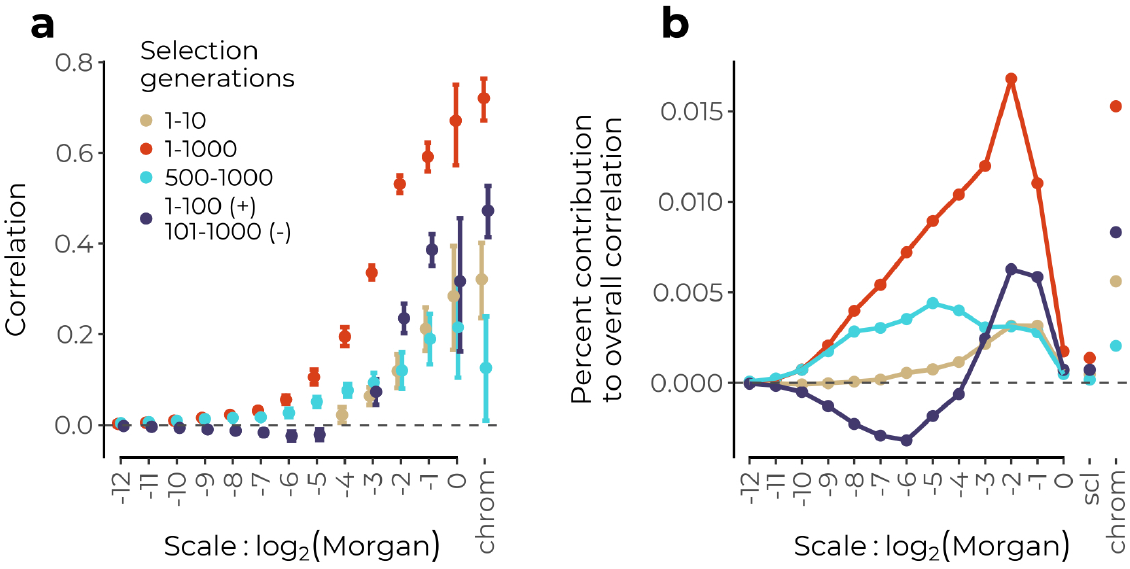
The wavelet decomposition of the correlation between recombination rate and introgressed ancestry can detect temporally-localized effects of selection in hybrids. **(a)** Selection acting only on the first 10 generations of recombinant hybrids generates significant positive wavelet correlations only at broad scales (brown) (viewed in generation 1000), whereas continuous selection over 1000 generations continues to generate correlations on finer scales (red). Selection that begins after generation 500 and of neutral mixture and continues until generation 1000 also generates fine-scale correlations, as well as broad-scale correlations. When selection acts continuously but reverses direction after 100 generations to favor the alternate ancestry, positive broad-scale correlations persist as negative correlations establish at finer scales. The comparison between these last two cases indicates that correlations at broad scales are dominated by the dynamics of selection in the early generations after mixture. Only significant correlations are shown, error bars represent 95% confidence intervals across 20 replicate simulations. **(b)** The correlations in **(a)** are weighted by the variance at each scale to give the contribution of each scale to the total correlation.

**Figure S8:**
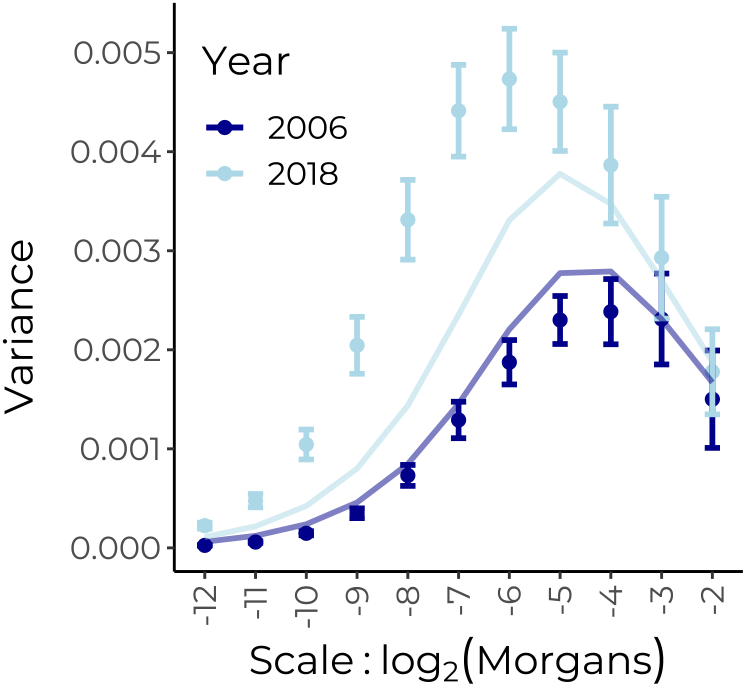
Power spectrum of proportion of malinche-like tracts in a hybrid swordtail population. Points show the empirical power spectra of proportion of malinche-like tracts in the two samples at the ends of the start and end of the time series. Error bars are 95% confidence intervals from a weighted block jacknife across chromosomes. Lines show theoretical expectations under neutrality, assuming 114 generations since initial admixture in the 2018 sample (previously estimate, Schumer lab), a generation time of 3 generations/year, and hybrid population size of 500 diploids, and an admixture proportion of 0.3.

**Figure S9:**
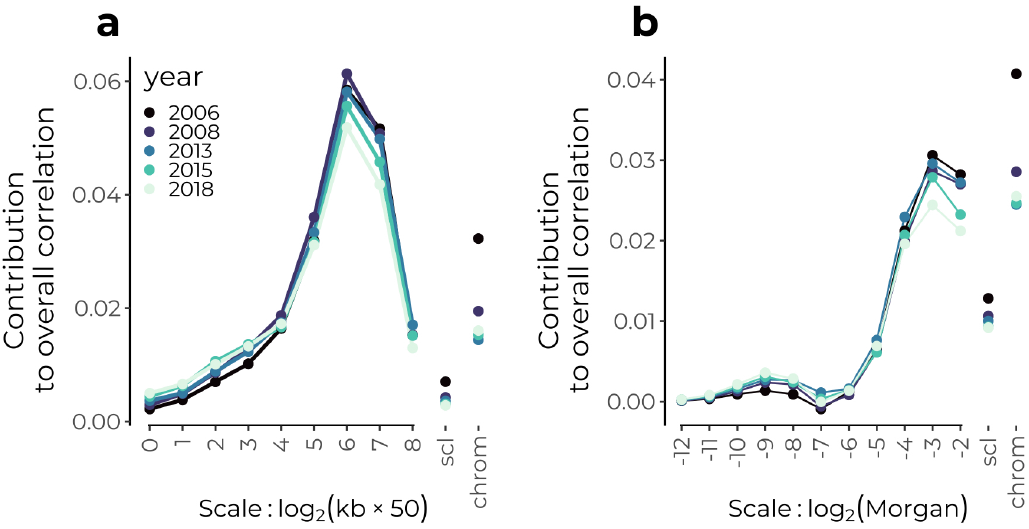
Contribution of each genomic scale to overall correlation between recombination and malinche-like ancestry. Results shown on the physical map **(a)** and genetic map **(b)**.

**Figure S10:**
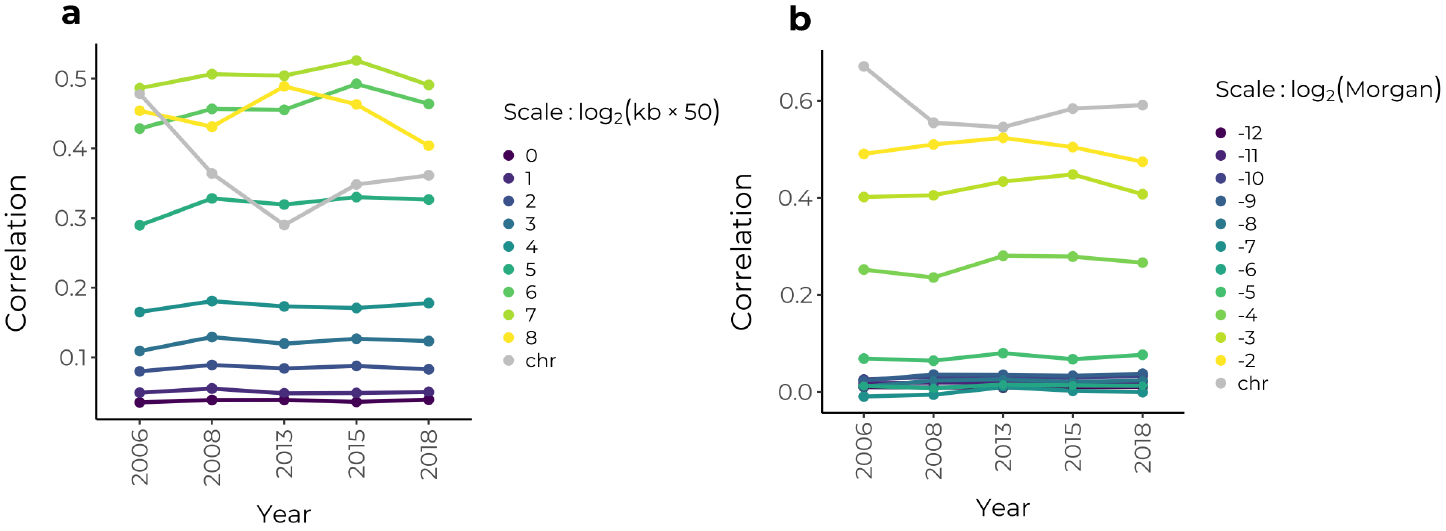
Correlations through time at varying spatial scales between recombination rate and minor parent ancestry (*X. malinche*). Results shown on the physical map **(a)** and genetic map **(b)**. Correlations between are not significantly different between any two timepoints at any scale.

**Figure S11:**
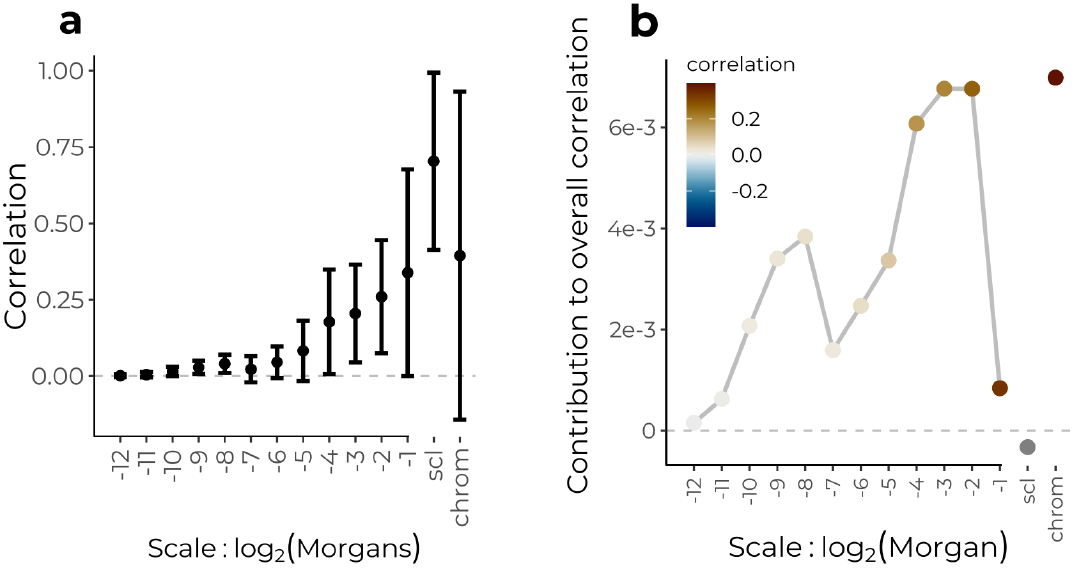
**(a)** Correlations across scales between recombination rate and anubis-like ancestry on the genetic map. **(b)** Contribution of each scale to the overall correlation (Eqn. 3).

**Figure S12:**
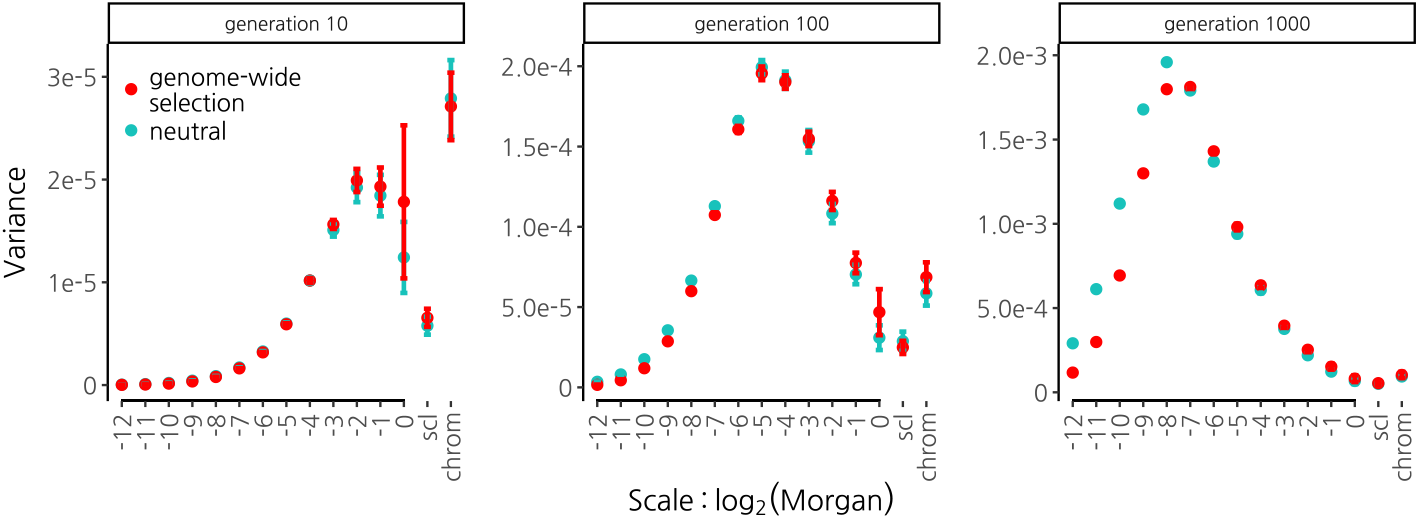
Power spectra of ancestry proportion for simulations with widespread weak selection (red) vs. neutral simulations (blue). Selection acts against alleles from one ancestry with additive effects within and across loci at 10,000 loci. Each allele has a selection coefficient of −2 × 10^−5^, yielding a 20% fitness reduction in F1s. Error bars are 95% confidence intervals computed from 20 replicate simulations.

**Figure S13:**
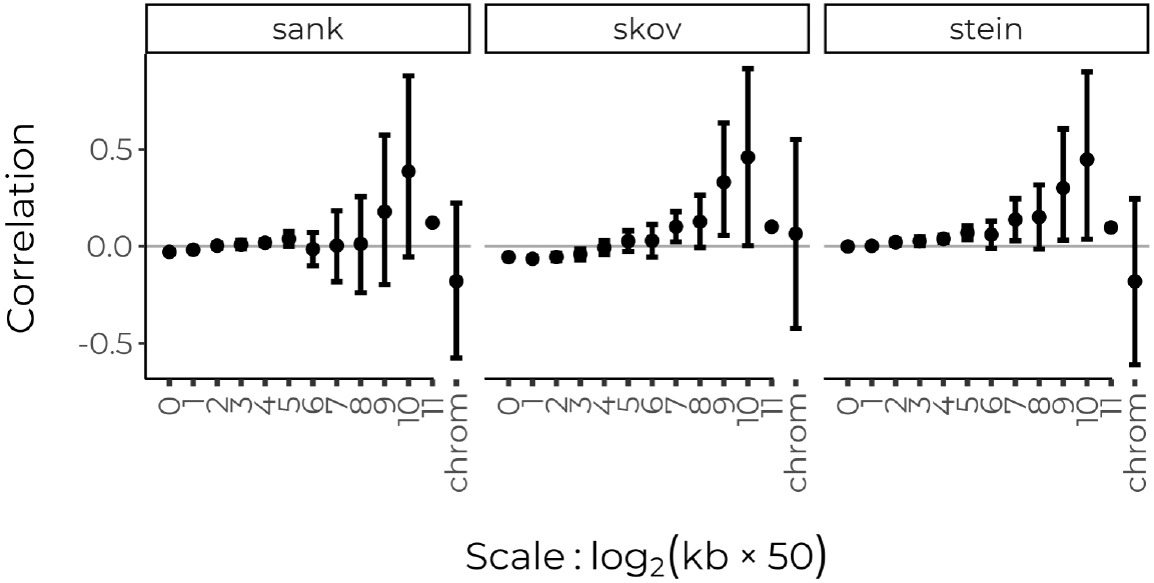
Correlations across scales between recombination rate and proportion of Neanderthal-like haplotypes on the human autosomes. **Left)** Data from Sankararaman *et al*. (2014) for the CEU sample of 1000 genomes. **Middle)** Data from Skov *et al*. (2020). **Right)** Data from Steinrücken *et al*. (2018) for the CEU population. Recombination rates are from Halldorsson *et al*. (2019) and are log10-transformed. Error bars show 95% confidence intervals from a weighted jackknife over chromosomes.

**Figure S14:**
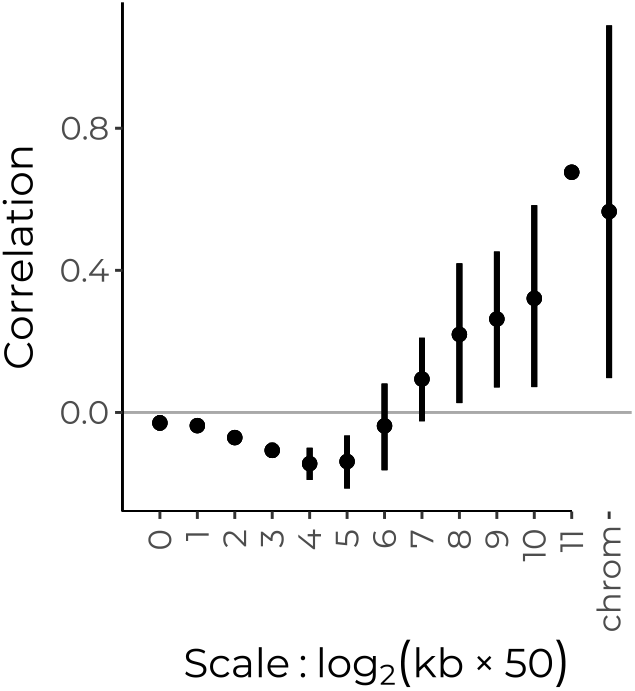
Correlations across scales between recombination rate and density of coding sequence base pairs on the physical map of the human autosomes.

**Figure S15:**
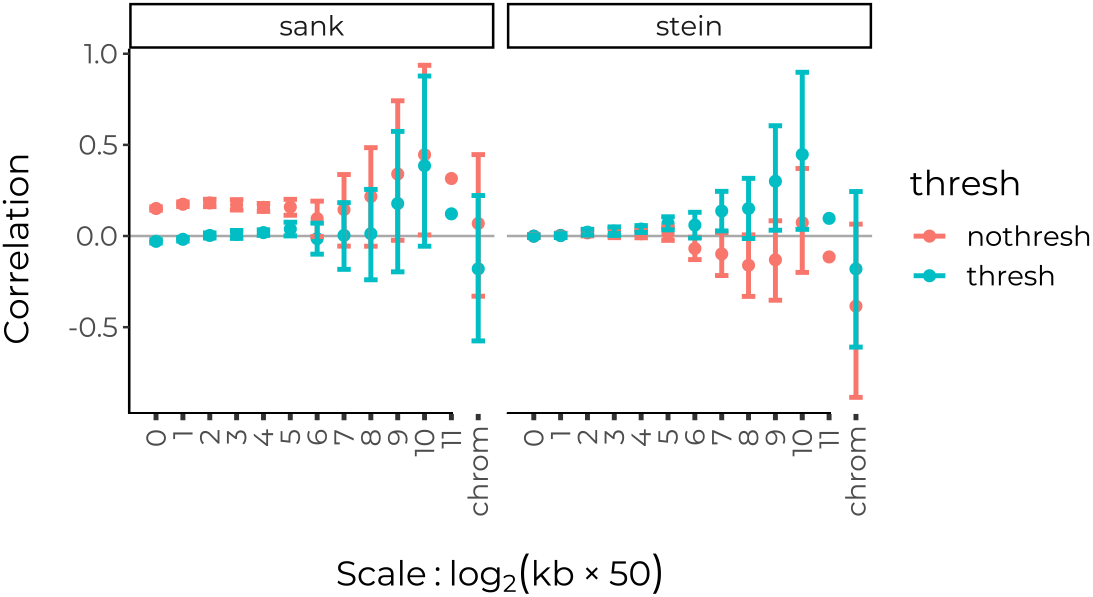
Effect of posterior threshold on correlations across scales between recombination rate and sample proportion of Neanderthal-like haplotypes estimated by Sankararaman *et al*. (2014) (left) and Steinrücken *et al*. (2018) (right). Following the methods described in these papers, we applied a thresholds of 0.9 and 0.42, respectively, to the posterior probabilities of a site matching a Neanderthal reference. We note that two studies that are commonly cited for evidence of selection against Neanderthal alleles utilize the non-thresholded posterior values from this study - Figure 2 in Sankararaman *et al*. (2014) and the work of Juric *et al*. (2016).

## Notes

### Competing Interest Statement

The authors have declared no competing interest.

### Summary of Updates

Corrected spelling of first author's name, in the meta data. Added italics to the abstract.

## References

Aeschbacher S, Selby JP, Willis JH, Coop G. 2017. Population-genomic inference of the strength and timing of selection against gene flow. Proceedings of the National Academy of Sciences. 114:7061–7066.

Barton N, Bengtsson BO. 1986. The barrier to genetic exchange between hybridising populations. Heredity. 57:357–376.

Barton NH. 1983. Multilocus clines. Evolution. 37:454–471.

Berner D, Roesti M. 2017. Genomics of adaptive divergence with chromosome-scale heterogeneity in crossover rate. Molecular ecology. 26:6351–6369.

Calfee E, Gates D, Lorant A, Perkins MT, Coop G, Ross-Ibarra J. 2021. Selective sorting of ancestral introgression in maize and teosinte along an elevational cline. PLoS genetics. 17:e1009810.

Chan AH, Jenkins PA, Song YS. 2012. Genome-wide fine-scale recombination rate variation in Drosophila melanogaster. PLOS Genetics. 8:e1003090.

Coop G. 2022. Genetic similarity and genetic ancestry groups. arXiv preprint arXiv:2207.11595. .

Coyne JA, Orr HA. 2004. Speciation. Sinauer Associates, Sunderland, MA.

Edelman NB, Frandsen PB, Miyagi M, Clavijo B, Davey J, Dikow RB, García-Accinelli G, Van Belleghem SM, Patterson N, Neafsey DE et al. 2019. Genomic architecture and introgression shape a butterfly radiation. Science. 366:594–599.

Gravel S. 2012. Population genetics models of local ancestry. Genetics. 191:607–619.

Green RE, Krause J, Briggs AW, Maricic T, Stenzel U, Kircher M, Patterson N, Li H, Zhai W, Fritz MHY et al. 2010. A draft sequence of the Neandertal genome. Science. 328:710–722.

Hajdinjak M, Mafessoni F, Skov L, Vernot B, Hübner A, Fu Q, Essel E, Nagel S, Nickel B, Richter J et al. 2021. Initial Upper Palaeolithic humans in Europe had recent Neanderthal ancestry. Nature. 592:253–257.

Halldorsson BV, Palsson G, Stefansson OA, Jonsson H, Hardarson MT, Eggertsson HP, Gunnarsson B, Oddsson A, Halldorsson GH, Zink F et al. 2019. Characterizing mutagenic effects of recombination through a sequence-level genetic map. Science. 363:eaau1043.

Haller BC, Galloway J, Kelleher J, Messer PW, Ralph PL. 2019. Tree-sequence recording in SLiM opens new horizons for forward-time simulation of whole genomes. Molecular ecology resources. 19:552–566.

Haller BC, Messer PW. 2019. Slim 3: forward genetic simulations beyond the wright–fisher model. Molecular biology and evolution. 36:632–637.

Hamid I, Korunes KL, Beleza S, Goldberg A. 2021. Rapid adaptation to malaria facilitated by admixture in the human population of Cabo Verde. Elife. 10:e63177.

Harris K, Nielsen R. 2016. The Genetic Cost of Neanderthal Introgression. Genetics. 203:881–891.

Harrison RG, Larson EL. 2014. Hybridization, introgression, and the nature of species boundaries. Journal of Heredity. 105:795–809.

Juric I, Aeschbacher S, Coop G. 2016. The strength of selection against Neanderthal introgression. PLOS Genetics. 12:e1006340. Publisher: Public Library of Science.

Kim BY, Huber CD, Lohmueller KE. 2018. Deleterious variation shapes the genomic landscape of introgression. PLoS Genetics. 14:e1007741.

Liang M, Nielsen R. 2014. The lengths of admixture tracts. Genetics. 197:953–967.

Loh PR, Lipson M, Patterson N, Moorjani P, Pickrell JK, Reich D, Berger B. 2013. Inferring admixture histories of human populations using linkage disequilibrium. Genetics. 193:1233–1254.

Maples BK, Gravel S, Kenny EE, Bustamante CD. 2013. Rfmix: a discriminative modeling approach for rapid and robust local-ancestry inference. The American Journal of Human Genetics. 93:278–288.

Martin SH, Davey JW, Salazar C, Jiggins CD. 2019. Recombination rate variation shapes barriers to introgression across butterfly genomes. PLOS Biology. 17:e2006288.

Maynard Smith J, Haigh J. 1974. The hitch-hiking effect of a favourable gene. Genetics Research. 23:23–35.

Moran BM, Payne C, Langdon Q, Powell DL, Brandvain Y, Schumer M. 2021. The genomic consequences of hybridization. eLife. 10:e69016.

Moran BM, Payne CY, Powell DL, Iverson EN, Banerjee SM, Madero A, Gunn TR, Langdon QK, Liu F, Matney R et al. 2022. A lethal genetic incompatibility between naturally hybridizing species in Mitochon-drial Complex I. BioRxiv. pp. 2021–07.

Nachman MW, Payseur BA. 2012. Recombination rate variation and speciation: theoretical predictions and empirical results from rabbits and mice. Philosophical Transactions of the Royal Society B: Biological Sciences. 367:409–421. Publisher: Royal Society.

National Academies of Sciences, Engineering, and Medicine. 2023. Using population descriptors in genetics and genomics research: A new framework for an evolving field. .

Nosil P, Funk DJ, Ortiz-Barrientos D. 2009. Divergent selection and heterogeneous genomic divergence. Molecular ecology. 18:375–402.

Percival DB, Walden AT. 2000. Wavelet methods for time series analysis. volume 4. Cambridge University Press.

Powell DL, García-Olazábal M, Keegan M, Reilly P, Du K, Díaz-Loyo AP, Banerjee S, Blakkan D, Reich D, Andolfatto P et al. 2020. Natural hybridization reveals incompatible alleles that cause melanoma in swordtail fish. Science. 368:731–736.

Powell DL, Payne C, Banerjee SM, Keegan M, Bashkirova E, Cui R, Andolfatto P, Rosenthal GG, Schumer M. 2021. The genetic architecture of variation in the sexually selected sword ornament and its evolution in hybrid populations. Current Biology. 31:923–935.

Price AL, Tandon A, Patterson N, Barnes KC, Rafaels N, Ruczinski I, Beaty TH, Mathias R, Reich D, Myers S. 2009. Sensitive Detection of Chromosomal Segments of Distinct Ancestry in Admixed Populations. PLOS Genetics. 5:e1000519.

Pugach I, Matveyev R, Wollstein A, Kayser M, Stoneking M. 2011. Dating the age of admixture via wavelet transform analysis of genome-wide data. Genome Biology. 12:R19.

Racimo F, Sankararaman S, Nielsen R, Huerta-Sánchez E. 2015. Evidence for archaic adaptive introgression in humans. Nature Reviews Genetics. 16:359–371.

Ravinet M, Faria R, Butlin RK, Galindo J, Bierne N, Rafajlović M, Noor MaF, Mehlig B, Westram AM. 2017. Interpreting the genomic landscape of speciation: a road map for finding barriers to gene flow. Journal of Evolutionary Biology. 30:1450–1477.

Rieseberg LH, Whitton J, Gardner K. 1999. Hybrid zones and the genetic architecture of a barrier to gene flow between two sunflower species. Genetics. 152:713–727.

Rogers J, Mahaney MC, Witte SM, Nair S, Newman D, Wedel S, Rodriguez LA, Rice KS, Slifer SH, Perelygin A et al. 2000. A genetic linkage map of the baboon (papio hamadryas) genome based on human microsatellite polymorphisms. Genomics. 67:237–247.

Samuels A, Altmann J. 1986. Immigration of a Papio anubis male into a group of Papio cynocephalus baboons and evidence for an anubis-cynocephalus hybrid zone in amboseli, kenya. International Journal of Primatology. 7:131–138.

Samuk K, Owens GL, Delmore KE, Miller SE, Rennison DJ, Schluter D. 2017. Gene flow and selection interact to promote adaptive divergence in regions of low recombination. Molecular ecology. 26:4378–4390.

Sanderson J, Sudoyo H, Karafet TM, Hammer MF, Cox MP. 2015. Reconstructing Past Admixture Processes from Local Genomic Ancestry Using Wavelet Transformation. Genetics. 200:469–481.

Sankararaman S, Mallick S, Dannemann M, Prüfer K, Kelso J, Pääbo S, Patterson N, Reich D. 2014. The landscape of Neandertal ancestry in present-day humans. Nature. 507:354–357.

Sankararaman S, Sridhar S, Kimmel G, Halperin E. 2008. Estimating local ancestry in admixed populations. The American Journal of Human Genetics. 82:290–303.

Schumer M, Xu C, Powell DL, Durvasula A, Skov L, Holland C, Blazier JC, Sankararaman S, Andolfatto P, Rosenthal GG et al. 2018. Natural selection interacts with recombination to shape the evolution of hybrid genomes. Science. 360:656–660.

Skov L, Coll Maciá M, Sveinbjörnsson G, Mafessoni F, Lucotte EA, Einarsdóttir MS, Jonsson H, Halldorsson B, Gudbjartsson DF, Helgason A et al. 2020. The nature of Neanderthal introgression revealed by 27,566 Icelandic genomes. Nature. 582:78–83.

Skov L, Hui R, Shchur V, Hobolth A, Scally A, Schierup MH, Durbin R. 2018. Detecting archaic introgression using an unadmixed outgroup. PLoS Genetics. 14:e1007641.

Spencer CCA, Deloukas P, Hunt S, Mullikin J, Myers S, Silverman B, Donnelly P, Bentley D, McVean G. 2006. The influence of recombination on human genetic diversity. PLOS Genetics. 2:e148.

Steinrücken M, Spence JP, Kamm JA, Wieczorek E, Song YS. 2018. Model-based detection and analysis of introgressed Neanderthal ancestry in modern humans. Molecular ecology. 27:3873–3888.

Tang H, Choudhry S, Mei R, Morgan M, Rodriguez-Cintron W, Burchard EG, Risch NJ. 2007. Recent genetic selection in the ancestral admixture of puerto ricans. The American Journal of Human Genetics. 81:626–633.

Telis N, Aguilar R, Harris K. 2020. Selection against archaic hominin genetic variation in regulatory regions. Nature Ecology & Evolution. 4:1558–1566.

Tung J, Charpentier MJ, Garfield DA, Altmann J, Alberts SC. 2008. Genetic evidence reveals temporal change in hybridization patterns in a wild baboon population. Molecular ecology. 17:1998–2011.

Veller C, Edelman NB, Muralidhar P, Nowak MA. 2023. Recombination and selection against introgressed DNA. Evolution. 77:1131–1144.

Vernot B, Akey JM. 2014. Resurrecting surviving Neandertal lineages from modern human genomes. Science. 343:1017–1021.

Vilgalys TP, Fogel AS, Anderson JA, Mututua RS, Warutere JK, Siodi IL, Kim SY, Voyles TN, Robinson JA, Wall JD et al. 2022. Selection against admixture and gene regulatory divergence in a long-term primate field study. Science. 377:635–641.

Whitcher B. 2022. waveslim: Basic Wavelet Routines for One-, Two-, and Three-Dimensional Signal Processing. R package version 1.8.4.

Wu CI. 2001. The genic view of the process of speciation. Journal of evolutionary biology. 14:851–865.

Yair S, Lee KM, Coop G. 2021. The timing of human adaptation from Neanderthal introgression. Genetics. 218:iyab052.

